# Myeloid FtH Regulates Macrophage Response to Kidney Injury by Modulating Snca and Ferroptosis

**DOI:** 10.1101/2025.03.25.645219

**Authors:** Tanima Chatterjee, Sarah Machado, Kellen Cowen, Mary Miller, Yanfeng Zhang, Laura Volpicelli-Daley, Lauren Fielding, Rudradip Pattanayak, Frida Rosenblum, László Potor, György Balla, József Balla, Christian Faul, Abolfazl Zarjou

## Abstract

This study explored the role of myeloid ferritin heavy chain (FtH) in coordinating kidney iron trafficking in health and disease. Synuclein-α (Snca) was the sole iron-binding protein upregulated in response to myeloid FtH deletion (FtH^Δ/Δ^). Following kidney injury, FtH^Δ/Δ^ mice showed worsened kidney function. Transcriptome analysis revealed coupling of FtH deficiency with ferroptosis activation, a regulated cell death associated with iron accumulation. Adverse effects of ferroptosis were evidenced by upregulation of ferroptosis-related genes, increased oxidative stress markers, and significant iron deposition in kidney tissues. This iron buildup in FtH^Δ/Δ^ kidneys stemmed from macrophage reprogramming into an iron-recycling phenotype, driven by Spic induction. Mechanistically, we establish that monomeric Snca functions as a ferrireductase catalyst, intensifying oxidative stress and triggering ferroptosis. Additionally, Snca accumulates in kidney diseases distinguished by leukocyte expansion across species. These findings position myeloid FtH as a pivotal orchestrator of the FtH-Snca-Spic axis driving macrophage reprogramming and kidney injury.

**Highlights:**

- Myeloid FtH deficiency drives kidney injury via activation of ferroptosis
- MΦ FtH deficiency induces Snca, linking iron dysregulation to MΦ function and response to kidney injury
- Ferrireductase activity of monomeric Snca augments oxidative stress, promoting lipid peroxidation and ferroptosis

**In brief:** MΦ FtH modulates Snca and Spic to coordinate the injury response, linking iron trafficking to ferroptosis-induced kidney injury

## INTRODUCTION

Acute kidney injury (AKI) is a prevalent clinical challenge contributing to considerable morbidity and affecting approximately one-quarter of hospitalized patients, with no currently available effective therapeutic interventions.^1,2^ Following AKI, excessive tissue damage and maladaptive repair mechanisms are well-recognized drivers of chronic kidney disease (CKD) development and progression.^3–5^ CKD is detrimental as it increases cardiovascular risk, reduces quality of life and life expectancy, leading to high mortality and morbidity rates while imposing a considerable financial burden on healthcare systems.^6,7^ Mechanistically, myeloid cells, particularly macrophages (MΦ), play a pivotal role in orchestrating the response to tubular damage and facilitating repair.^8,9^

Despite the diverse etiologies of AKI, many exhibit an immune component involving MΦ.^10–12^ MΦ comprise a unique, and heterogeneous population with remarkable phenotypic plasticity, enabling them to respond dynamically to a wide array of environmental signals. The extent of MΦ expansion observed in preclinical models and human biopsies directly correlates with the severity of kidney injury.^12,13^ Moreover, MΦ are integral to maintaining iron homeostasis, a process increasingly recognized as central in kidney pathology.^14–16^

Iron is essential for aerobic life; however, when unrestrained, it catalyzes the formation of reactive oxygen species (ROS), leading to cellular and tissue injury and triggering a regulated form of cell death known as ferroptosis.^17,18^ Consequently, balancing efficient iron distribution with mitigation of its potential toxicity is critical. Evolution has addressed this challenge with ferritin, a large intracellular nanocage capable of safely storing up to 4500 iron atoms.^19^ Ferritin is a spherical protein complex composed of heavy (FtH) and light (FtL) chains. Through its ferroxidase activity, FtH converts the pro-oxidant ferrous (Fe^2+^) iron into the less reactive ferric (Fe^3+^) form, thereby facilitating safe storage.^19,20^ The indispensable role of FtH is underscored by the early embryonic lethality observed in mice with global FtH deletion, highlighting the paramount role of its ferroxidase activity.^21^

Given their versatility and critical role in iron metabolism, MΦ are increasingly recognized as therapeutic targets across various disease settings to regulate iron availability, mitigate organ damage, and promote recovery.^22^ Building on this insight, we investigated the effects of myeloid-specific FtH deficiency on iron trafficking in kidney health and disease. We found upregulation of synuclein-⍰ (Snca) in MΦ in response to FtH deletion. Snca is a small, intrinsically disordered protein primarily known for its role in neurodegenerative diseases, particularly Parkinson’s disease.^23,24^ However, emerging evidence indicates that Snca also contributes to immune regulation and possesses pro-inflammatory properties.^25,26^ Snca is expressed in diverse cell types, with its highest expression in monocytes among peripheral leukocytes.^27,28^ Additionally, Snca’s expression and function are directly linked to iron metabolism. It is able to bind metals including iron, a process that facilitates its accumulation, and in a self-propagating manner, accelerates tissue damage.^29–31^ Moreover, monomeric Snca possesses ferrireductase activity, potentially driving iron-mediated oxidative stress via generation of pro-oxidant ferrous iron.^32,33^ In this context, Snca appears to play a dual role, contributing to immune regulation by promoting a proinflammatory microenvironment while also influencing iron metabolism. This positions Snca as a potentially key factor in inflammatory and degenerative diseases beyond the nervous system. However, the pathogenic role of MΦ Snca expression in the context of kidney disease, and its interaction with FtH and iron, is not known and constitutes the focus of this report.

## RESULTS

### Myeloid FtH deletion triggers Snca induction in macrophages

To explore the transcriptional alterations linked to FtH deletion in myeloid cells, we conducted an unbiased bulk RNA-sequencing analysis of the kidneys from wildtype (FtH^fl/fl^) and myeloid specific FtH deficient (FtH^Δ/Δ^) mice under quiescent conditions. We identified multiple significantly modulated genes, with Snca notably upregulated in the kidneys of FtH^Δ/Δ^ mice compared to controls (Figure 1A). Among genes that encode proteins with iron binding capacity, only Snca revealed significant upregulation in FtH^Δ/Δ^ kidneys when compared to FtH^fl/fl^ controls (Figures 1B, S1A). Immunostaining for Snca in the kidneys revealed a similar tubular staining pattern among genotypes, most prominent in outer cortical stripe (Figure S1B). In contrast, a higher number of Snca-positive interstitial cells were observed in FtH^Δ/Δ^ kidneys compared to FtH^fl/fl^ controls, indicating elevated expression in tissue-resident MΦ (Figure 1C, S1B). This observation was further supported by the marked Snca expression in the splenic red pulp and interstitial cells resembling hepatic Kupffer cells in FtH^Δ/Δ^ livers (Figure 1C). To validate the monocytic lineage source of Snca signal, we performed immunofluorescence double staining on kidneys, spleens, and livers of FtH^fl/fl^, and FtH^Δ/Δ^ mice by targeting Snca and CD11b (Figure 1D). Neutrophils were ruled out as the source of Snca upregulation using Ly6G immunostaining (Figure S2.) Furthermore, using a publicly available transcriptomic database, we confirmed dominant expression of Snca in monocytic cell lineage when compared to other leukocytes (Figure S3). Using the spleen as a rich source of monocytes, we validated and quantified significant induction of monomeric Snca in response to FtH deletion (Figure 1E-G). Consistent with its characterization as a secretory protein,^34–36^ we found that MΦ upregulation of Snca was also associated with a substantial increase in serum levels in FtH^Δ/Δ^ mice (Figure 1H). These findings suggest that FtH deficiency drives aberrant Snca overexpression and accumulation within MΦ in various tissues including the kidneys.

**Figure 1.**
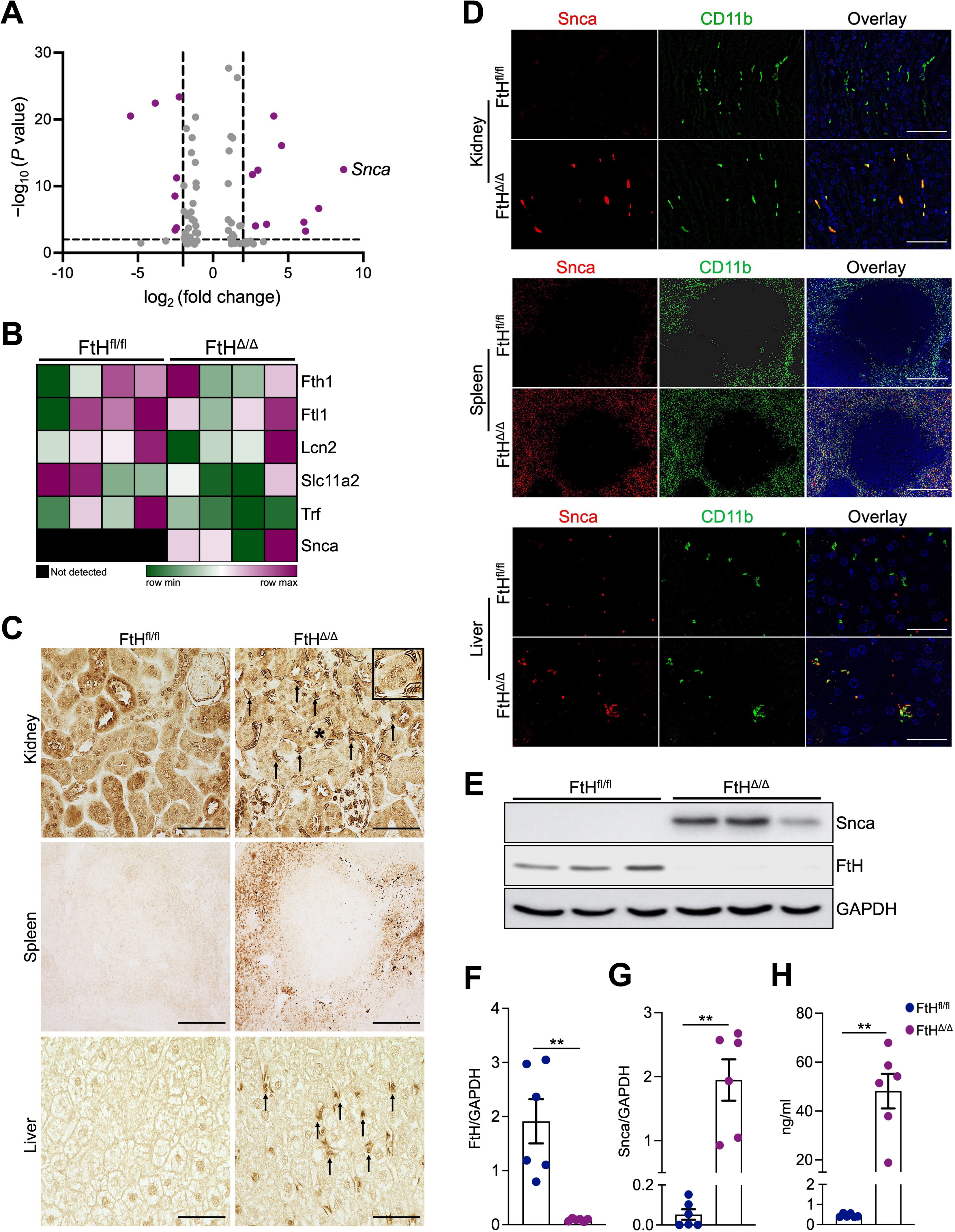
Myeloid FtH deletion triggers Snca induction in macrophages. (A) Volcano plot showing differentially expressed genes in the kidneys of wildtype (FtH^fl/fl^) and myeloid-specific FtH deficient (FtH^Δ/Δ^) mice under quiescent conditions following bulk-RNA sequencing (n=4/group). (B) Heatmap illustrating the expression of genes that encode iron-binding proteins, Fth1, Ftl1, Lcn2 (Lipocalin-2), Slc11a2 (divalent metal transporter-1), Trf (transferrin), in kidneys of FtH^fl/fl^ and FtH^Δ/Δ^ mice using normalized reads obtained via bulk-RNA sequencing. (C) Representative immunohistochemistry using an anti-Snca antibody in the kidney, spleen, and liver of FtH^fl/fl^ and FtH^Δ/Δ^ mice under homeostatic conditions. Black arrows in the FtH^Δ/Δ^ kidney indicate Snca expressing interstitial cells. Inset: higher magnification of a tubule with surrounding Snca expressing cells, marked with an asterisk. Scale bars: kidney= 100 μm, spleen= 200 μm, liver= 50 μm. (D) Immunofluorescence staining of Snca and the myeloid marker CD11b in the kidney, spleen, and liver of FtH^fl/fl^ and FtH^Δ/Δ^ mice under baseline conditions. Scale bars: kidney= 25 μm, spleen= 200 μm, liver= 25 μm. (E) Representative western blot of FtH and Snca expression levels in FtH^fl/fl^ and FtH^Δ/Δ^ spleen at baseline. GAPDH was used as a loading control. (F, G) Densitometric analysis of spleen FtH and Snca expression, normalized to GAPDH (n=6/genotype). (H) Serum Snca levels measured by ELISA (n=6/genotype). **P < 0.01.

### Myeloid FtH deficiency exacerbates AKI and accelerates AKI-to-CKD progression

To elucidate the functional significance of myeloid FtH in kidney disease, we employed a well-established and readily adjustable aristolochic acid (AA)-induced model of AKI-to-CKD transition, characterized by pronounced tubular toxicity, robust inflammation, and substantial leukocyte accumulation. Mice from both genotypes received vehicle or AA and kidney function was monitored via serial serum creatinine measurements at weeks 1, 3, and 6 post final AA injection (Figure 2A). Following the initial rise, wildtype mice showed a gradual decline in serum creatinine levels, although these values never fully returned to baseline (Figure 2B). In contrast, FtH^Δ/Δ^ mice exhibited significantly elevated serum creatinine levels at all time points compared to FtH^fl/fl^ controls, indicating that myeloid FtH deficiency leads to unresolved kidney injury. (Figure 2B). Heightened CKD progression in FtH^Δ/Δ^ mice was further characterized by increased proteinuria, coupled with higher urinary Snca excretion, and augmented collagen deposition (Figure 2C-F). Next, we evaluated Snca expression in kidney MΦ at six weeks post-injury and found that nearly all CD11b⁺ cells from both genotypes expressed Snca. This robust induction was also evident in wildtype animals, although the signal was markedly more pronounced in FtH^Δ/Δ^ kidneys (Figure 2G). Congruent with these results, western blot analysis revealed that AA-induced kidney injury led to increased Snca levels in both FtH^fl/fl^ and FtH^Δ/Δ^ kidneys, with FtH^Δ/Δ^ mice exhibiting markedly higher accumulation (Figure 2H). Histological analysis of kidney sections at week 6 post-AA injection revealed substantial leukocyte expansion in both genotypes, which was strikingly higher in FtH^Δ/Δ^ mice (Figure 2I). To determine the cellular basis of the intensified leukocyte expansion in FtH^Δ/Δ^ kidneys, we conducted flow cytometric analysis of kidney leukocyte populations (Figure 2J). This analysis revealed that the augmented leukocyte counts in FtH^Δ/Δ^ kidneys following AA-induced injury were primarily driven by the expansion of MΦ, and neutrophils (Figure 2H). Taken together, these findings demonstrate that myeloid FtH deletion exacerbates kidney injury, impairs repair processes, promotes excessive leukocyte expansion, and increases Snca deposition, culminating in accelerating kidney disease progression.

**Figure 2.**
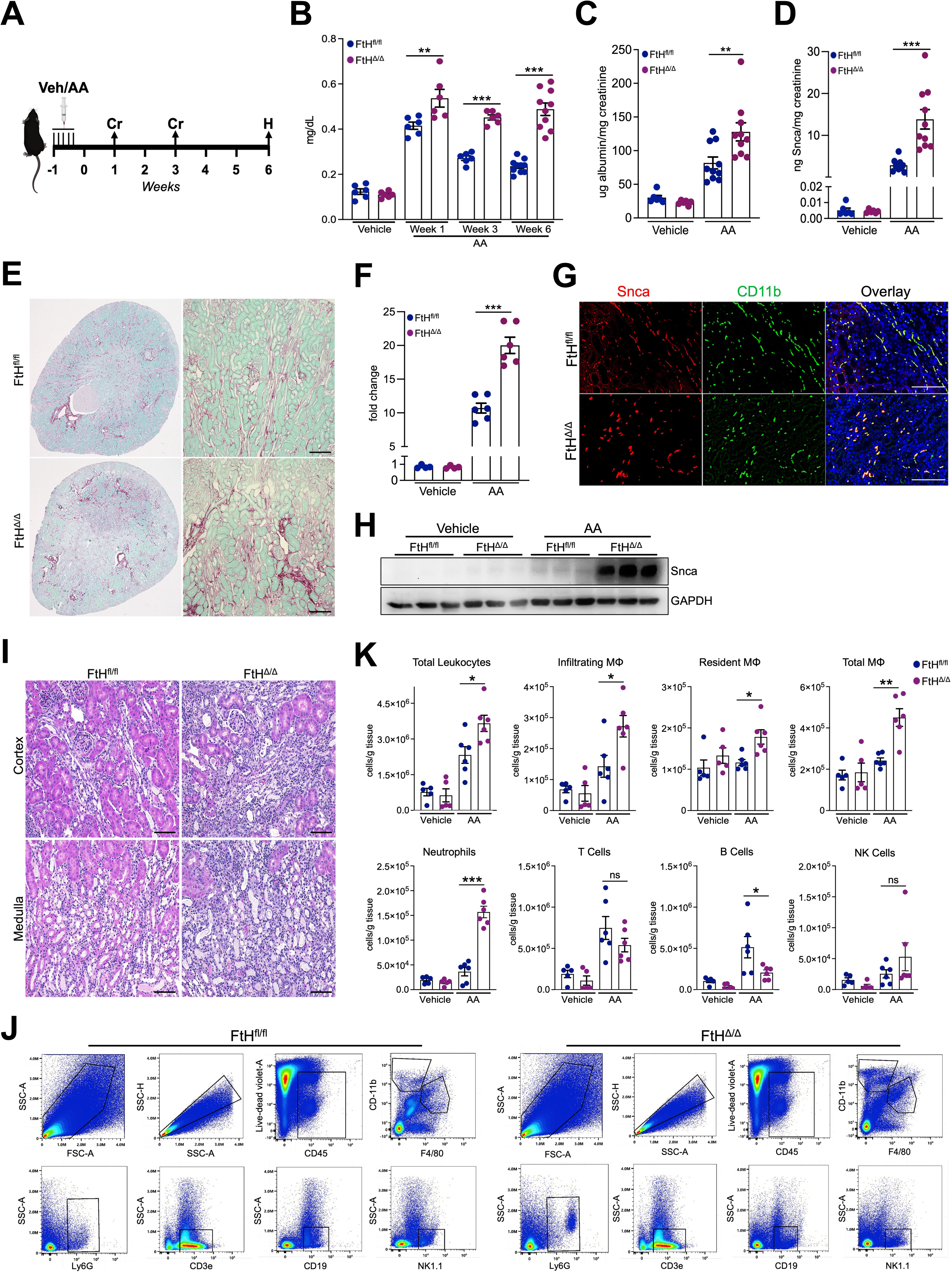
Myeloid FtH deficiency exacerbates AKI and accelerates AKI-to-CKD progression. (A) Schematic representation of the experimental design. Mice were administered aristolochic acid (AA) or vehicle, with measurements and analyses performed at indicated time points. Cr: creatinine measurement; H: Harvest and endpoint analyses. (B) Serum creatinine levels were measured at indicated time points in FtH^fl/fl^ and FtH^Δ/Δ^ mice (Vehicle, n=6/genotype, AA, n=6/genotype for week1, and week3 AA, n=10/genotype for week6 AA). (C) Albumin was measured in the urine of vehicle and AA treated mice and normalized to creatinine (Vehicle, n=6/genotype; AA, n=10/genotype). (D) Snca was measured in the urine of vehicle and AA treated animals and normalized to creatinine (Vehicle, n=6/genotype; AA, n=10/genotype). (E) Representative sirius red/fast green stain demonstrating collagen deposition at 6 weeks post AA administration. Scale bar = 200 μm. (F) Quantification of kidney fibrosis (fold change) based on sirius red/fast green stain in FtH^fl/fl^ and FtH^Δ/Δ^ mice using Image J (Vehicle, n=4/genotype; AA, n=6/genotype). (G) Representative immunofluorescence staining of kidney sections at six weeks post-AA induced injury analyzed for Snca and CD11b, with DAPI nuclear counterstaining. Scale bar = 25 μm. (H) Representative western blot of monomeric Snca expression in kidney lysates from vehicle and AA treated mice at six weeks. (I) Representative H&E stain of FtH^fl/fl^ and FtH^Δ/Δ^ kidneys at week six following AA induced CKD. Scale bar = 100 μm. (J) Representative flow cytometry histograms of kidney immune cell populations in FtH^fl/fl^ and FtH^Δ/Δ^ mice at six weeks post AA administration (Vehicle, n=5/genotype; AA, n=6/genotype). Representative gating strategy for identifying leukocytes (CD45^+^), macrophages (F4/80^+^), neutrophils (Ly6G^+^), T cells (CD3e^+^), B cells (CD19^+^), and NK cells (NK1.1^+^). (K) Quantification of immune cell populations in kidney tissues at six weeks post vehicle or AA administration (Vehicle, n=5/genotype; AA, n=6/genotype). Data are represented as number of cells per gram kidney tissue. ns = not significant, *P < 0.05, **P < 0.01, ***P < 0.001.

### Single-cell transcriptomic analysis of kidney leukocytes reveals ferroptosis induction following FtH deletion

To examine the transcriptional response to FtH deletion, we performed single-cell RNA sequencing (ScRNA-seq) on kidney leukocytes isolated from FtH^fl/fl^ and FtH^Δ/Δ^ mice six weeks after AA or vehicle treatment (Figure 3A). CD45⁺ immune cells were sorted from kidneys and subjected to scRNA-seq using the 10× Genomics platform. Unbiased clustering of single-cell transcriptomic data identified nine distinct immune cell populations (Figure 3B, S4). Top differentially expressed genes (DEGs) analysis confirmed the presence of all major immune cell types, including monocytes/MΦ, dendritic cells (DCs), neutrophils, innate lymphoid cells (ILCs), B cells, and three distinct T-cell clusters (Figure 3B, C). KEGG pathway analysis of FtH-deficient cells revealed ferroptosis as the most significantly activated pathway, both under quiescent conditions and following AA administration (Figure 3D, E). Additionally, pathways associated with protein processing, chemokine signaling, and antigen presentation were enriched, highlighting a dysregulated immune response in the absence of FtH (Figure 3D). Heatmap of differentially enriched genes in FtH deficient cells compared to wildtype counterparts, and dotplot of individual genes engaged in ferroptosis are demonstrated in Figure S5. Given that Lyz2-Cre is expressed in immune cells beyond the monocytic lineage, we conducted a gene set enrichment analysis (GSEA) specifically on monocytes/MΦ to evaluate the enrichment of the glutathione metabolism pathway, rapidly induced during ferroptosis, as well as the ferroptosis pathway itself. This analysis confirmed the activation of both pathways in FtH-deficient monocytes/MΦ, regardless of injury state (Figure 3F, G). Additionally, the dot plot shows a robust induction of key ferroptosis-related genes in MΦ following FtH deletion, observed under both vehicle and AA treatment conditions (Figure 3H). These results infer that loss of FtH increases susceptibility of MΦ to iron metabolism perturbations and subsequent activation of ferroptosis in the kidney.

**Figure 3.**
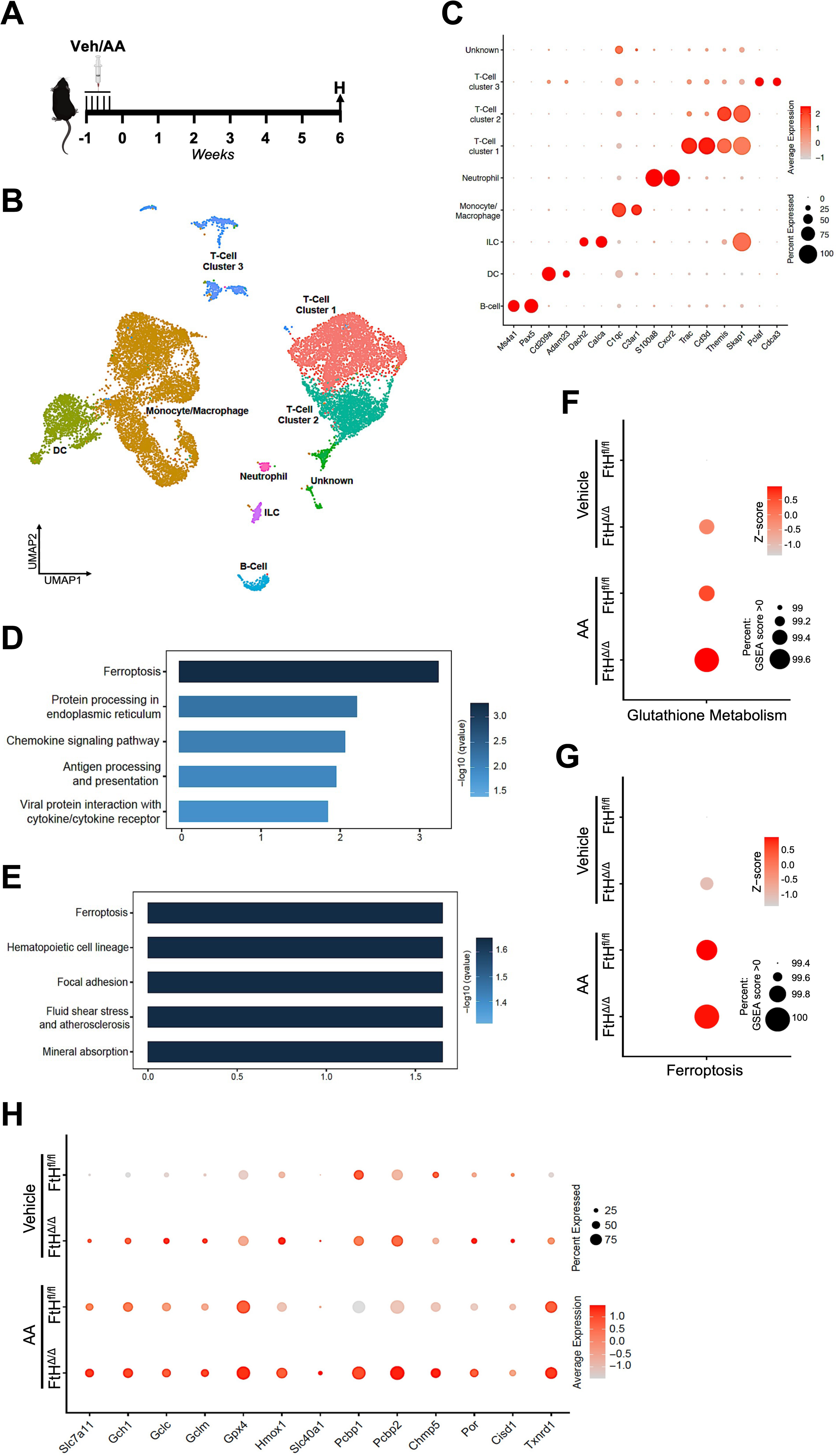
Single-cell transcriptomic analysis of leukocytes reveals ferroptosis induction following FtH deletion. (A) Schematic of the experimental set up. FtH^fl/fl^ and FtH^Δ/Δ^ mice received vehicle or AA five consecutive days. Mice were harvested for single cell RNA sequencing analysis of kidney leukocytes six weeks after AA administration. H indicates the harvest and endpoint analysis. (B) Uniform manifold approximation and projection (UMAP) plot showing the clustering of immune cells (CD45^+^) in the kidney based on single-cell RNA sequencing. The analysis identifies nine distinct clusters, with contaminating kidney cells and clusters representing less than 1% removed. (C) Cell type-specific expression of marker genes for manually annotated clusters. Dot size denotes percentage of cells expressing the marker. Color scale represents average gene expression values. (D) Pathway enrichment analysis of differentially expressed genes in FtH deficient cells versus wildtype cells under vehicle-treated conditions. (E) Pathway enrichment analysis of differentially expressed genes in FtH deficient cells versus wildtype cells following AA administration. (F-G) Gene set enrichment analysis (GSEA) for (F) glutathione metabolism and (G) ferroptosis pathways in monocytes/MΦ from vehicle- and AA-treated FtH^fl/fl^ and FtH^Δ/Δ^ mice. The size of the dots represents the percentage of genes enriched in the pathway, while the color indicates the Z-score. (H) Dot plot showing key genes involved in ferroptosis across different genotypes and experimental conditions. Dot size denotes percentage of cells expressing the marker. Color scale represents average gene expression values.

### Ferroptosis drives exacerbated kidney injury in myeloid FtH-deficient mice

Given the wave-like propagation of ferroptosis within affected tissues, we reasoned that the exacerbated injury and CKD progression observed in FtH^Δ/Δ^ kidneys may be driven by increased ferroptotic activity. This deduction was examined by assessing lipid peroxidation and ferroptosis-associated marker levels. Western blot analysis revealed significantly elevated levels of 4-hydroxynonenal (HNE), a byproduct of lipid peroxidation, and acyl-CoA synthetase long-chain family member 4 (ACSL4), a key regulator of ferroptosis, in FtH^Δ/Δ^ kidneys compared to FtH^fl/fl^ controls (Figure 4A-D). We also examined these patterns in kidneys under baseline conditions and while a trend toward higher levels in FtH^Δ/Δ^ kidneys was observed, the difference did not reach statistical significance (Figure S6). To further investigate the ferroptotic signaling cascade, we quantified the mRNA expression of other key regulatory genes. In FtH^Δ/Δ^ mice, nuclear factor erythroid 2-related factor 2 (Nrf2), a central antioxidant regulator rapidly activated during ferroptotic injury, was significantly upregulated post injury (Figure 4E). Similarly, Slc7a11 and Slc40a1 (ferroportin), which are critical for glutathione metabolism and iron export respectively, were markedly upregulated (Figure 4F, G). To assess the functional impact of ferroptosis in this model, we administered ferrostatin-1 (Fer-1), a potent and specific ferroptosis inhibitor, at specific intervals before, during, and after AA-induced kidney injury (Figure 4H). Notably, Fer-1 treatment effectively reduced serum creatinine levels in FtH^Δ/Δ^ mice to levels comparable with FtH^fl/fl^ controls treated with AA alone or in combination with Fer-1, highlighting ferroptosis as the primary driver of the excessive injury observed in FtH^Δ/Δ^ mice (Figure 4I). Concordantly, collagen deposition and the degree of albuminuria were comparable between both genotypes, with no significant differences observed following AA and Fer-1 treatment (Figure 4K-L). Collectively, these findings indicate that ferroptosis plays a key role in the heightened kidney injury observed in FtH^Δ/Δ^ mice.

**Figure 4.**
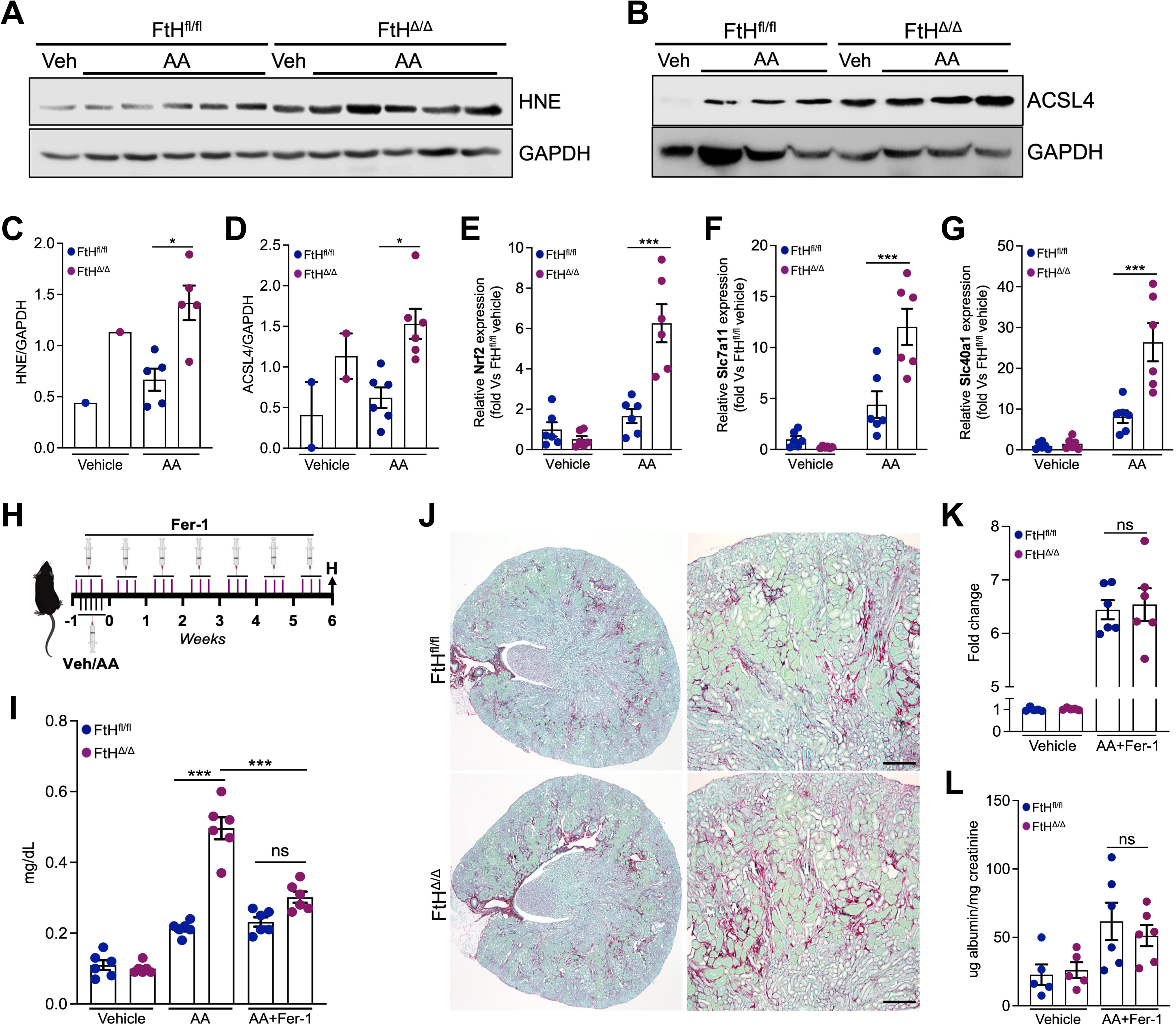
Ferroptosis drives exacerbated kidney injury in myeloid FtH**^Δ^**^/**Δ**^ mice. (A, B) Representative western blot showing (A) HNE and (B) ACSL4 expression levels in kidney lysates from mice treated with vehicle or AA. (C) Densitometric quantification of HNE expression normalized to GAPDH (Vehicle, n=1/genotype; AA, n=5/genotype). (D) Densitometric quantification of HNE expression normalized to GAPDH (Vehicle, n=1/genotype; AA, n=6/genotype). (E-G) QRT-PCR analysis of mRNA expression levels of (E) Nrf2, (F) Slc7a11, and (G) Slc40a1 in kidneys at six weeks post treatment with vehicle or AA. Data are normalized to GAPDH and represented as fold vs FtH^fl/fl^ vehicle group (Vehicle, n=6/genotype; AA, n=6/genotype). (H) Schematic illustrating the experimental design for ferrostatin-1 (Fer-1) treatment to diminish ferroptosis induced injury. (I) Serum creatinine levels across genotypes, and experimental conditions at six weeks following indicated treatment (Vehicle, n=6/genotype; AA, n=6/genotype). (J) Representative sirius red/fast green staining demonstrating collagen deposition at 6 weeks post AA+Fer-1 administration in FtH^fl/fl^ and FtH^Δ/Δ^ mice (Vehicle, n=6/genotype; AA+Fer-1, n=6/genotype), Scale bar = 200 μm. (K) Quantification of kidney fibrosis (fold change) based on sirius red/fast green stain in FtH^fl/fl^ and FtH^Δ/Δ^ mice post AA+Fer-1 administration using ImageJ (Vehicle, n=6/genotype; AA+Fer-1, n=6/genotype). (L) Urinary albumin levels measured in vehicle and AA+Fer-1 treated mice and normalized to urine creatinine (Vehicle, n=6/genotype; AA+Fer-1, n=6/genotype). ns = not significant, *P < 0.05, ***P < 0.001.

### FtH deficiency promotes macrophage transition to an iron-recycling phenotype via Spic upregulation, enhancing iron accumulation

Next, using modified Perl’s stain to enhance histochemical detection of iron, we assessed iron accumulation, a hallmark of ferroptosis, in the injured kidneys of FtH^fl/fl^ and FtH^Δ/Δ^ mice and observed more prominent iron deposition in FtH^Δ/Δ^ kidneys at 6 weeks post-AA treatment (Figure 5A). We postulated that a disrupted crosstalk between MΦ and tubular cells in FtH^Δ/Δ^ kidneys could contribute to their heightened susceptibility to abnormal iron deposition. To test this premise, we utilized a previously established model of parenteral iron overload as illustrated in Figure 5B. Three days after the final injection, kidney sections stained with Prussian blue revealed distinctly increased iron deposition in FtH^Δ/Δ^ mice compared to FtH^fl/fl^ controls (Figure 5C). This finding was further validated by assessing FtH and FtL protein levels as indicators of iron burden, which revealed a significant upregulation of both ferritin chains in kidneys of FtH^Δ/Δ^ mice (Figure 5D-F). These results were independent of circulating iron levels, as there were no genotype-dependent differences in serum iron levels in mice that received either vehicle or parenteral iron, despite a significant increase in serum iron following iron administration (Figure 5G). We then performed a metallomics analysis using inductively coupled plasma mass spectrometry (ICP-MS) to examine levels of iron, copper, zinc, and magnesium in kidneys, livers, and spleens of FtH^fl/fl^ and FtH^Δ/Δ^ mice following iron administration. We corroborated that FtH^Δ/Δ^ kidneys accumulated significantly higher levels of iron, while no significant differences were observed in the livers and spleens (Figure 5H). Levels of other studied metals were mostly similar and are presented in Figure S7. Collectively, these observations unravel a distinct propensity of FtH^Δ/Δ^ kidneys to iron accumulation. To explore the molecular underpinnings responsible for this process we examined the transcriptomic landscape of harvested kidneys following iron overload, using bulk-RNA sequencing. Volcano plot analysis revealed that Spic, the key transcription factor involved in the development of iron-recycling MΦ biological profile, was significantly upregulated in FtH^Δ/Δ^ kidneys (Figure 5I). Moreover, we observed a signature of genes associated with this phenotype to be significantly upregulated in FtH^Δ/Δ^ kidneys (Figure 5J, S8). Induction of Spic and Slc40a1, two critical genes associated with iron recycling phenotype, was independently verified through quantitative real time-PCR [qRT-PCR] (Figure 5K, L). These findings led us to postulate that the increased iron accumulation observed in FtH^Δ/Δ^ kidneys post-AA may also result from elevated Spic expression, indicating a higher number of iron recycling-MΦ. Indeed, our results show a marked induction of Spic in response to kidney injury in both genotypes, with a significantly higher induction in FtH^Δ/Δ^ kidneys (Figure 5M). In summary, these findings demonstrate that FtH deficient MΦ adopt an iron recycling profile distinguished by Spic upregulation and increased ferroportin (Slc40a1) expression, leading to augmented iron accumulation in the kidney.

**Figure 5.**
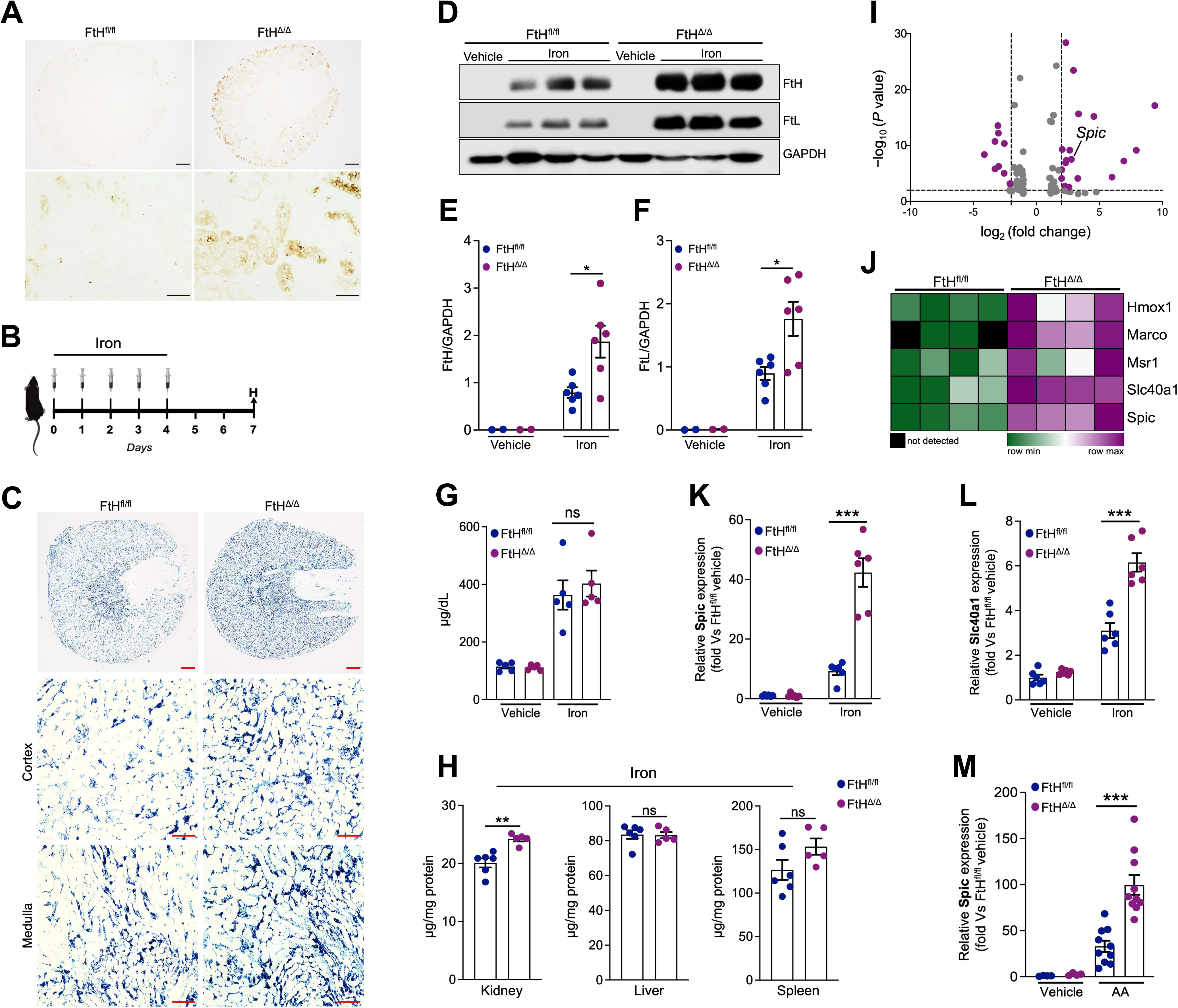
FtH deficiency promotes macrophage transition to an iron-recycling phenotype via Spic upregulation, enhancing iron accumulation. (A) Modified Perl’s Prussian blue staining demonstrating iron deposition in kidneys of FtH^fl/fl^ and FtH^Δ/Δ^ mice at six weeks post AA treatment. Scale bars: 200 μm (top), 50 μm (bottom). (B) Schematic of the iron overload model. (C) Perls’ Prussian blue staining of kidney sections from iron treated mice. Lower panels highlight iron accumulation in cortex and medulla. Scale bars: 200 μm (top), 100 μm (middle and bottom rows). (D) Representative western blot analysis of FtH and FtL protein expression in kidneys of mice treated with vehicle or iron. (E, F) Densitometric analysis of (E) FtH and (F) FtL protein levels, normalized to GAPDH (Vehicle, n=2/genotype; Iron, n=6/genotype). (G) Serum iron levels of mice treated with vehicle or iron (Vehicle, n=5/genotype; Iron, n=5/genotype). (H) Tissue iron content in kidneys, livers, and spleens of iron treated FtH^fl/fl^ and FtH^Δ/Δ^ mice was measured using ICP-MS (FtH^fl/fl^, n=6; FtH^Δ/Δ^, n=5). (I) Volcano plot of significantly modulated genes following iron administration as determined by bulk-RNA sequencing, n=4 per group. (J) Heatmap of genes linked to iron recycling profile of MΦ in response to iron loading in kidneys of FtH^fl/fl^ and FtH^Δ/Δ^ mice using normalized reads obtained via bulk-RNA sequencing. (K, L) QRT-PCR analysis of (K) Spic and (L) Slc40a1 mRNA expression in kidneys following treatment with vehicle or iron. Data are normalized to GAPDH and represented as fold vs FtH^fl/fl^ vehicle group (Vehicle, n=6/genotype; Iron, n=6/genotype). (M) Total RNA was isolated from kidneys at six weeks post vehicle or AA injection and analyzed for Spic expression. Data are normalized to GAPDH and represented as fold vs FtH^fl/fl^ vehicle group (Vehicle, n=6/genotype; AA, n=6/genotype). ns = not significant, *P < 0.05, ***P < 0.001.

### Monomeric Snca’s ferrireductase activity drives oxidative stress, triggering ferroptosis

To examine whether Snca can directly contribute to oxidative stress and ferroptosis, we cultured bone marrow-derived MΦ (BMDMs) from wildtype mice and treated them with increasing concentrations of recombinant monomeric Snca (0.1, 0.5, and 1 μM) for 16 hours. Western blot analysis confirmed a dose-dependent uptake of Snca by BMDMs (Figure 6A). Live-cell imaging revealed rapid Snca uptake that began within 20 minutes of incubation (Figure 6B). To assess Snca’s impact on lipid peroxidation, we used Western blot analysis to measure HNE and ACSL4 expression and demonstrate a significant increase in both markers following monomeric Snca treatment. (Figure 6C-E). Additionally, qRT-PCR analysis revealed upregulation of key ferroptosis-associated genes, Slc7a11 and Slc40a1, upon Snca exposure (Figure 6F, G). We subsequently examined the effect of Snca treatment on generation of intracellular ROS. Using DCFDA staining and flow cytometric analysis, we observed a dose-dependent increase in ROS-positive cells following Snca treatment compared to vehicle-treated cells (Figure 6H-J). Lipid peroxidation, a characteristic byproduct of ferroptosis, was assessed using the lipid peroxidation sensor BODIPY 581/591, and flow cytometric analysis. Notably, Snca treatment led to a dose-dependent increase in lipid peroxidation, closely aligning with the observed rise in ROS levels (Figure 6K-M). The dose-dependent increase in intracellular ROS burden following Snca treatment was further visualized through fluorescent microscopy (Figure 6N). To validate the paramount role of Snca’s ferrireductase activity in ROS generation, we used three approaches to diminish this activity. These included using boiled or proteinase-K treated Snca, as well as aggregated (Snca fibrils) Snca, which loses its ferrireductase function due to improper protein folding. Western blot analysis demonstrated that only native monomeric Snca, but not its aggregated or denatured forms, induced HNE accumulation (Figure 6O-P). Together, these results indicate that monomeric Snca promotes oxidative stress via its ferrireductase activity, contributing to lipid peroxidation and ferroptosis in BMDMs.

**Figure 6.**
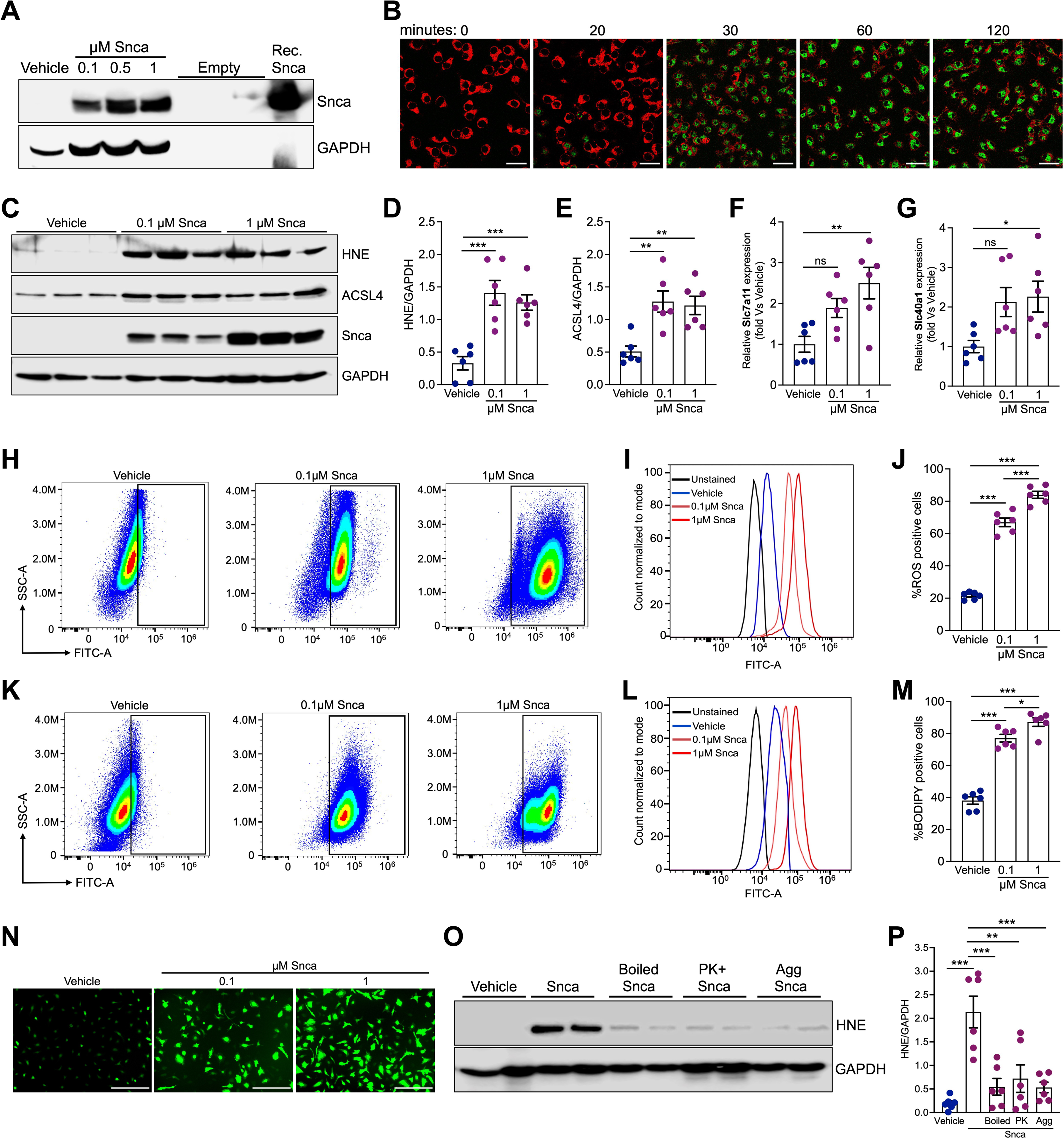
Monomeric Snca’s ferrireductase activity drives oxidative stress: A hallmark of ferroptosis. (A) Representative Western blot showing dose-dependent uptake of recombinant monomeric Snca (0.1, 0.5, and 1 μM) by BMDMs after 16 hours of incubation. Recombinant Snca was used as a positive control. (B) For uptake assays, BMDMs were treated with A488-labeled recombinant Snca, and the A488 signal was monitored using live-cell imaging over a time course of 0–120 minutes. The green signal indicates successful internalization of the recombinant monomeric Snca. Scale bar = 100 μm. (C) Representative Western blot showing HNE and ACSL4 expression levels in BMDMs treated with vehicle (PBS) or Snca (0.1 and 1 μM). (D-E) Densitometric quantification of (D) HNE and (E) ACSL4 expression, normalized to GAPDH (Vehicle, n=6; Snca 0.1 μM, n=6; Snca 1 μM, n=6). (F-G) QRT-PCR analysis of mRNA expression levels of (F) Slc7a11 and (G) Slc40a1 in BMDMs treated with vehicle or Snca (0.1 and 1 μM). Data are normalized to GAPDH and represented as fold vs vehicle group (Vehicle, n=6; Snca 0.1 μM, n=6; Snca 1 μM, n=6). (H) Representative flow cytometry plot showing ROS detection using DCFDA in BMDMs treated with vehicle or Snca (0.1 μM and 1 μM). The x-axis represents fluorescence intensity (indicative of ROS levels), while the y-axis represents cell count. (I) Representative histogram of DCFDA fluorescence intensity. (J) Quantification of ROS-positive cells following Snca treatment (Vehicle, n=6; Snca 0.1 μM, n=6; Snca 1 μM, n=6). (K) Representative flow cytometry plot showing lipid peroxidation levels measured using BODIPY 581/591 staining in BMDMs treated with vehicle or Snca (0.1 μM and 1 μM). The x-axis represents fluorescence intensity (indicative of lipid peroxidation levels), while the y-axis represents cell count. (L) Representative histogram of BODIPY fluorescence intensity. (M) Quantification of BODIPY positive cells (Vehicle, n=6; Snca 0.1 μM, n=6; Snca 1 μM, n=6). (N) Representative images demonstrating intracellular ROS using DCFDA in BMDMs treated with increasing concentrations of Snca (0.1 μM and 1 μM) compared to vehicle treated controls. Scale bar = 100 μm. (O) Representative Western blot demonstrates HNE expression in BMDMs treated with vehicle, monomeric Snca, boiled monomeric Snca, proteinase K-treated monomeric Snca, or aggregated Snca (fibrils) for 16 hours. (P) Densitometric quantification of HNE expression, normalized to GAPDH (Vehicle, n=6; boiled monomeric Snca 1 μM, n=6; proteinase K-treated monomeric Snca 1 μM, n=6; aggregated Snca (fibrils) 1 μM, n=6). ns = not significant, *P < 0.05, **P < 0.01, ***P < 0.001.

### Snca accumulation is a hallmark of kidney diseases characterized by leukocyte expansion across species

To explore the dynamics of Snca accumulation and its correlation with various kidney disease settings, we employed four distinct models: ischemia-reperfusion (I/R) injury, AA-and cisplatin-induced nephropathy, and a mouse model of Alport syndrome (Col4a3^−/−^) with spontaneous progression of CKD. Western blot analysis revealed a significant increase in Snca protein levels in the kidneys of mice subjected to AA-induced nephropathy at 6 weeks compared to vehicle treated controls (Figure 7A). Similarly, in the I/R injury model, Snca levels progressively increased over time, with the most pronounced upregulation observed at days 7 and 28 post-injury (Figure 7B). This pattern suggests that Snca may play a critical role in the transition from AKI-to-CKD, with its expression increasing over time rather than during the initial acute phase. In contrast, kidneys from cisplatin-treated mice and mice that mimic Alport syndrome did not show accumulation of Snca (Figure 7C, D). Densitometric quantification confirmed a significant upregulation of Snca in both the AA and I/R models (Figure 7E, F). Conversely, Snca expression was reduced in the cisplatin model, which is characterized by tubular toxicity and leukopenia, while no change was observed in the Alport mouse model (Figure 7G, H). Kidney injury of all four models was confirmed by serum creatinine measurements (Figure 2B, 7I, 7J).

**Figure 7.**
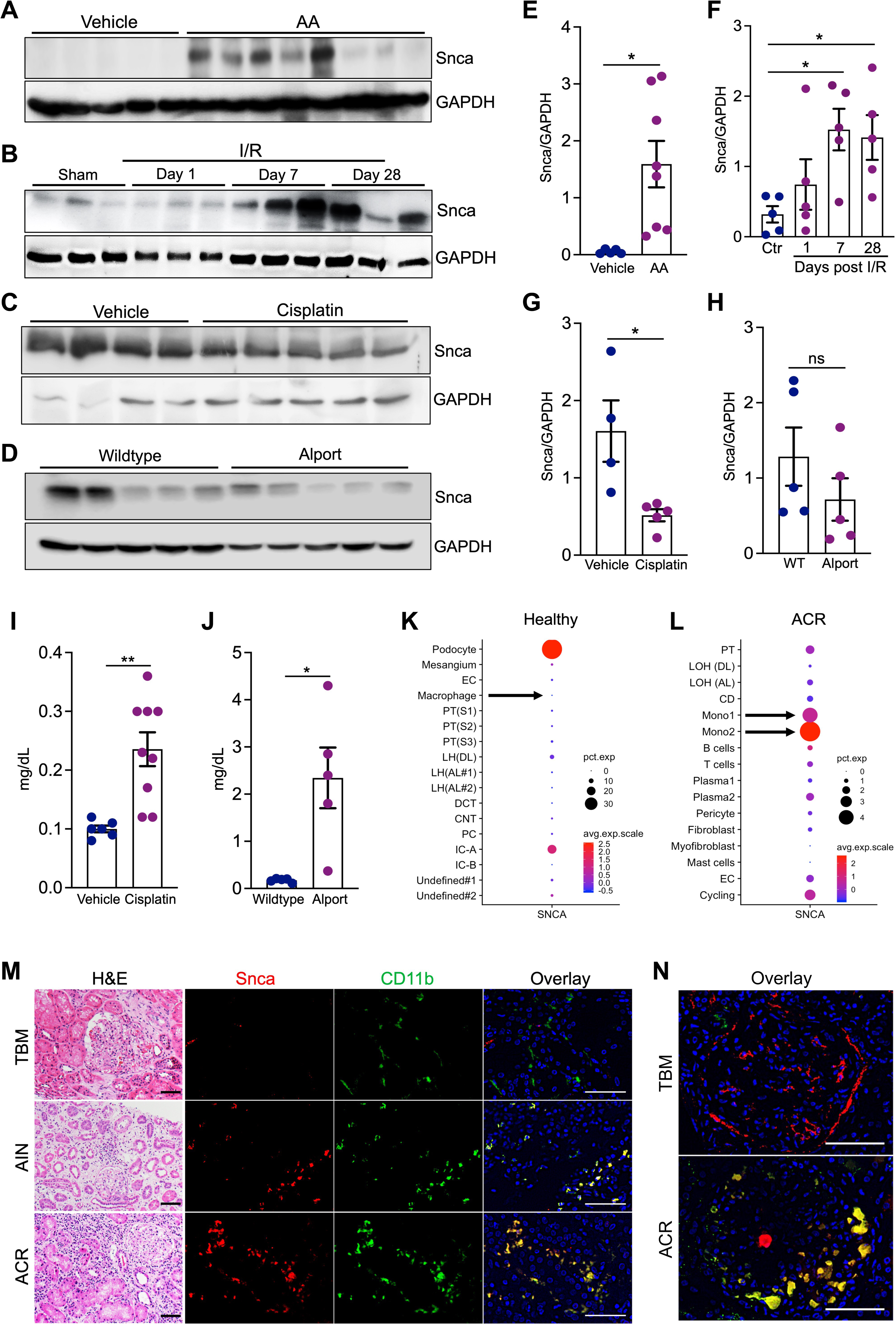
Snca accumulation is a hallmark of kidney diseases marked by leukocyte expansion across species. (A-D) Western blot analysis of monomeric Snca in kidney lysates from four different mouse models of kidney disease: (A) Wildtype mice were treated with vehicle or aristolochic acid (AA), and kidney lysates were collected six weeks post-AA administration. (B) Wildtype mice subjected to 20 minutes of ischemia-reperfusion (I/R) injury. Kidney samples were collected at indicated time-points and analyzed for Snca expression. (C) Protein expression of Snca in wildtype kidneys treated with cisplatin as described in methods. (D) Snca expression in kidneys from wildtype (Col4a3^+/+^) and Alport (Col4a3^−/−^) mice at twelve weeks of age. (E-H) Densitometric analysis of Snca protein expression in nephropathy models: (E) AA (Vehicle, n=5; AA, n=8), (F) I/R (Sham, n=5; day 1, 7, 28, n=5/timepoint). (G) cisplatin (Vehicle, n=4; cisplatin, n=5), and (H) Alport syndrome (n=5/genotype), normalized to GAPDH. (I) Serum creatinine levels in vehicle and cisplatin treated mice (Vehicle, n=6; cisplatin, n=9). (J) Serum creatinine in wildtype and Alport mice measured at 12 weeks of age (n=5/genotype). (K, L) Analysis of Snca expression using publicly available ScRNA-seq datasets from healthy (K) and (L) rejecting human kidney allograft (ACR). Arrows indicate monocyte/MΦ expression of Snca in (K) healthy and (L) ACR kidneys. (M) H&E stain (left panels) highlights histopathological features of human kidney biopsies from patients with thin basement membrane (TBM), acute interstitial nephritis (AIN), and acute cellular allograft rejection (ACR). Immunofluorescence staining of Snca and CD11b validates overlap in AIN, and ACR, but not in TBM. Scale bars = H&E stains: 100 µm, immunofluorescence: 25 µm. (N) Higher magnification overlay images of a healthy appearing glomeruli in TBM and a case of glomerulitis in ACR. Scale bars: 25 µm. ns = not significant, *P < 0.05, **P < 0.01.

To explore the translational and clinical implications of MΦ Snca expression in human kidney disease, we analyzed publicly available snRNA-seq datasets from healthy human kidneys and rejecting kidney allografts. In healthy kidneys, Snca expression was predominantly restricted to podocytes, with minimal expression detected in MΦ (Figure 7K). However, in kidney allografts with acute cellular rejection (ACR) which is marked by heavy leukocytes expansion, Snca expression was markedly upregulated in monocytes, emerging as their primary source of expression (Figure 7L). These results align with our supplemental data, which demonstrate Snca enrichment in monocytic lineage cells during human AKI (Figure S9, Table S1, 2). In agreement with these findings, analysis of human diabetic kidney disease, which is largely devoid of substantial leukocyte expansion, showed no significant Snca enrichment in leukocytes (Figure S10). These findings prompted us to further investigate the cellular localization of Snca in human kidney diseases with varying degrees of leukocyte involvement including thin basement membrane disease (TBN), where pathogenesis is largely independent of leukocyte expansion, and acute interstitial nephritis (AIN) and ACR, which are heavily leukocyte-dependent (Table S3). Routine histochemical, immunostaining, along with electron microscopy, was used by the pathologist to confirm the primary diagnosis (Figure 7M, S11). Immunofluorescence double staining revealed a lack of Snca expression in CD11b^+^ cells in TBM setting, whereas a pronounced upregulation of Snca colocalized with CD11b^+^ cells in diagnoses of AIN and ACR (Figure 7M). This finding was further reinforced by the presence of severe tubulitis and glomerulitis in ACR (Figure 7M, N). These results suggest that kidney diseases marked by leukocyte, particularly MΦ, expansion create a unique milieu that fosters Snca dysregulation, potentially amplifying the risk of Snca-mediated neurodegenerative diseases. This connection underscores the potential for kidney dysfunction to serve as a systemic driver of neurodegenerative processes, bridging inflammatory and iron-related pathologies across organ systems.

## DISCUSSION

In this study, we establish that deletion of FtH is associated with the upregulation of Snca in MΦ. Snca, known for its iron-binding and reducing properties, plays a critical role in the pathogenesis of several neurodegenerative diseases, including Parkinson’s disease.^23,24,31^ Following induction of AKI, FtH-deficient mice exhibited accelerated kidney disease progression accompanied by increased leukocyte expansion. Further analyses confirmed that activation of ferroptosis contributed to the injury observed in FtH^Δ/Δ^ mice. Mechanistically, our findings suggest that the ferrireductase activity of monomeric Snca promotes oxidative stress, thereby increasing lipid peroxidation and promoting ferroptosis. Moreover, we reveal that FtH deficiency triggers MΦ to adopt an iron-recycling phenotype through the upregulation of Spic, resulting in kidney iron accumulation, a hallmark of ferroptosis. Additionally, our findings indicate that monomeric Snca induction in MΦ is a defining feature of kidney diseases characterized by pronounced leukocyte aggregation in both mice and humans.

AKI and its progression to CKD is a multifactorial disorder that beyond its adverse impact on physiologic benchmarks, triggers maladaptive crosstalk between the kidney and other organs, exacerbating inflammation, oxidative stress, and cardiovascular pathology, ultimately driving high morbidity and mortality.^6,7,37^ Dysregulated iron metabolism is a common feature in AKI and is frequently observed during the clinical progression of CKD.^16,38–40^ Additionally, aberrant kidney iron deposition and ferroptosis have been identified as key drivers of AKI pathogenesis and the transition from AKI-to-CKD.^17,41,42^ First described in 2012,^43^ ferroptosis is a distinct and inherently iron dependent form of regulated cell death described by extensive iron accumulation and lipid peroxidation during the cell death process.^44^ Various ferroptosis-inducing factors can overwhelm the glutathione metabolism pathway, leading to an exhaustion of antioxidant capacity and ultimately an oxidative imbalance resulting in cellular damage and death.^44^ Interestingly, ferroptosis can propagate across cell populations in a synchronized wave-like fashion, leading to a characteristic spatiotemporal pattern of cell death.^45,46^ This process is likely intensified by increased Slc40a1 expression, which promotes the efflux of ferrous iron from cells undergoing ferroptosis, coupled with the release of iron-containing heme moieties from dying cells.^47^ Guided by our findings, we propose that FtH-deficient MΦ act as the primary drivers of ferroptosis in this model. This premise is supported by evidence demonstrating that ferroptosis is the most significantly enriched cellular pathway in FtH-deficient cells. This enrichment is accompanied by upregulation of markers associated with ferroptosis-mediated damage and activation of key ferroptosis regulators in whole kidneys. Importantly, experiments using Fer-1, a ferroptosis inhibitor,^48^ confirmed that the excess injury observed in FtH^Δ/Δ^ mice is primarily driven by ferroptosis.

Our findings indicate that an imbalance in the monocyte/MΦ iron recycling profile predisposes the kidneys of FtH^Δ/Δ^ mice to excessive iron deposition. Bulk RNA sequencing, supported by subsequent validation experiments, revealed a transcriptomic signature consistent with iron-recycling MΦ biological profile, a process coordinated by lineage defining transcription factor, Spic.^49^ Mammals have evolved an efficient strategy for iron recycling, where most of the required iron is retrieved from engulfed erythrocytes by a subset of MΦ known as iron-recycling MΦ. ^50,51^ Spic, a PU.1 related gene, is considered an indispensable transcription factor that selectively controls development of these highly specialized MΦ that express high levels of genes involved in iron trafficking including Slc40a1.^49,52–54^ Moreover, Spic^+^ MΦ have been proposed to engage in repair and resolution phase of injury.^55^ Our results suggest that the increased propensity of FtH deficient MΦ to upregulate Spic, along with the accompanying induction of Slc40a1, triggers the excessive iron accumulation observed in FtH^Δ/Δ^ kidneys under both iron overload and CKD conditions.

At the cellular level, regulation of key iron homeostasis proteins is orchestrated by iron regulatory proteins (IRPs) through their physical interaction with iron-responsive elements (IREs).^56,57^ These cytoplasmic IRPs bind with high affinity to conserved hairpin structures located in the untranslated regions of mRNAs (IREs), allowing cells to delicately balance the expression of iron import, storage, and export proteins in response to fluctuations in intracellular iron levels.^56,57^ Notably, the expression of Snca, similar to FtH, is controlled post-transcriptionally, as both mRNAs contain structured IREs in their 5′ untranslated regions that govern translation.^58,59^ Specifically, the Snca IRE is formed at the splice junction of the first two exons of the Snca gene.^60^ Based on existing evidence,^61,62^ and our findings, we propose that in the absence of FtH, an increase in the intracellular labile iron pool triggers the dissociation of IRPs from Snca mRNA, thereby enhancing its translation. However, contrary to ferroxidase activity of FtH that coverts the unstable ferrous form of iron into a more stable ferric form,^63^ Snca via its ferrireductase activity may augment oxidative stress by generating excess iron in ferrous state.^32^ While the exact function of neuronal Snca continues to be debated, its abundant presence and ferrireductase activity in red blood cells is proposed to play a crucial role in maintaining iron in its ferrous state within hemoglobin, which is essential for efficient oxygen binding.^64^ Similarly, within MΦ, Snca can contribute to generation of ROS, a fundamental process required for elimination of invading pathogens and coordinating signal transduction and cellular processes.^65^ Moreover, given that Slc40a1 exclusively exports iron in its ferrous state, it can be reasonably projected that Snca’s ferrireductase activity is essential for facilitating MΦ iron export. An expanding body of evidence increasingly substantiates the induction of Snca during inflammatory processes, highlighting its involvement in immune regulation.^28^ Moreover, Snca has been identified as a potent chemoattractant, a critical modulator of dendritic cell phenotypic maturation, and a pivotal immunomodulator, thus playing a central role in the orchestration of innate immunity and the maintenance of overall immune competence.^25,28,66–69^ Consistent with these findings, we observed an enrichment of inflammatory pathways in FtH-deficient monocytes under quiescent conditions, along with an increased leukocyte presence in the kidneys of FtH^Δ/Δ^ mice, which may, at least in part, be attributed to elevated Snca levels.

Due to its molecular weight of 14 kDa, Snca can traverse the glomerular filtration barrier and be excreted in the urine. Indeed, it was recently revealed that kidneys rapidly remove circulating Snca, a process that declines with worsening kidney disease, leading to Snca accumulation and ensuing aggregation into fibrils within the kidneys.^70^ These fibrils subsequently propagate to the brain, a process that is blocked by kidney denervation.^70^ While population studies indicate an elevated risk of Parkinson’s disease in patients with advanced CKD,^71–74^ the mechanisms that underpin this association remain unclear, particularly as most individuals with CKD do not develop Parkinson’s disease. This report offers a potential explanation for this discrepancy. We validate upregulation of Snca in MΦ across species in response to kidney injury. In mice, we demonstrate kidney accumulation of monomeric Snca in injury models characterized by predominant leukocyte expansion, such as I/R and AA. Conversely, kidney diseases with minimal leukocyte involvement in their pathogenesis, such as cisplatin-induced nephropathy and Col4a3 deficiency (Alport syndrome), do not exhibit increased Snca levels in kidneys. Similarly, while we observed increased presence of MΦ Snca in AIN and ACR, two kidney disease entities marked by high degree of leukocyte aggregation, such expression was absent in TBM disease. Moreover, through analysis of publicly available transcriptome datasets, we confirm that Snca expression in MΦ is minimal under both healthy conditions and diabetic injury but is notably elevated in ACR. Collectively, our findings suggest that the etiology of CKD, along with the extent of leukocyte (particularly MΦ) accumulation, may predispose the kidneys to Snca aggregation. This, in turn, could render this subgroup of patients more susceptible to subsequent brain accumulation of Snca via propagation from the kidneys.

In summary, our study uncovers a novel mechanistic pathway linking FtH deficiency in MΦ to enhanced kidney injury. We show that loss of FtH promotes the upregulation of Snca, an iron-binding, ferrireductase-active protein implicated in oxidative stress and ensuing ferroptosis, while simultaneously driving MΦ to adopt an iron-recycling profile via Spic upregulation. This dual effect leads to excessive iron accumulation and accelerated ferroptotic cell death in the kidney. Importantly, the elevated expression of Snca in MΦ correlates with kidney disease models characterized by marked leukocyte aggregation and may explain, at least in part, the observed association between CKD and an increased risk of neurodegenerative disorders. These findings highlight the intricate interplay between iron metabolism, FtH, Snca, and Spic in kidney injury and recovery, underscoring their collective influence on disease progression. By elucidating these mechanisms, our study provides a compelling rationale for targeting these pathways as potential therapeutic strategies to mitigate severity of AKI and by extension the process of AKI-to-CKD transition.

### Limitations of the study

Given the scope of the study, we did not investigate potential contribution of aggregated Snca in kidney disease development and progression. Moreover, future research is crucial to elucidate the mechanisms that make the kidneys, as opposed to the spleen or liver, particularly vulnerable to iron accumulation in response to myeloid FtH deficiency.

## RESOURCE AVAILABILITY

### Lead contact

Further information and requests for resources and reagents should be directed to and will be fulfilled by the lead contact, Abolfazl Zarjou

### Materials availability

This study did not generate new unique reagents. However, we welcome requests for clarification of protocols to ensure reproducibility of our findings.

### Data and code availability

The sequencing data for this study have been deposited at the Gene Expression Omnibus and are publicly available as of the date of publication. Accession numbers are listed in the key resources table. Any additional information required to reanalyze the data reported in this paper is available from the lead contact upon request. Please refer to supplemental information for the raw blot images.

## ACKNOWLEDGMENTS

We thank Dr. Anupam Agarwal for his critical insight and thorough review of this manuscript. This work was supported by National Institutes of Health (NIH) grant DK134402 (AZ), HUN-REN-DE (11003) and NKFIH ADVANCED 149734 (GyB, JB), and UAB-UCSD O’Brien Center for Acute Kidney Injury Research (NIH U54 DK137307).

## AUTHOR CONTRIBUTIONS

Conceptualization, T.C., J.B., Gy.B., C.F., L.V.D, A.Z.; methodology, T.C., S.M., K.C., M.M., L.P., R.P.; Investigation, T.C., Y.Z., M.M., K.C., R.P., L.P, A.Z; writing—original draft, T.C., S.M., A.Z.; writing—review & editing, T.C., S.M, K.C., M.M., Y.Z., F.R., L.P., J.B., Gy.B., L.V.D, C.F., A.Z.; funding acquisition, A.Z.; resources, F.R., J.B., L.P., L.V.D., L.F., A.Z.; supervision, A.Z.

## STAR⍰METHODS

### KEY RESOURCES TABLE

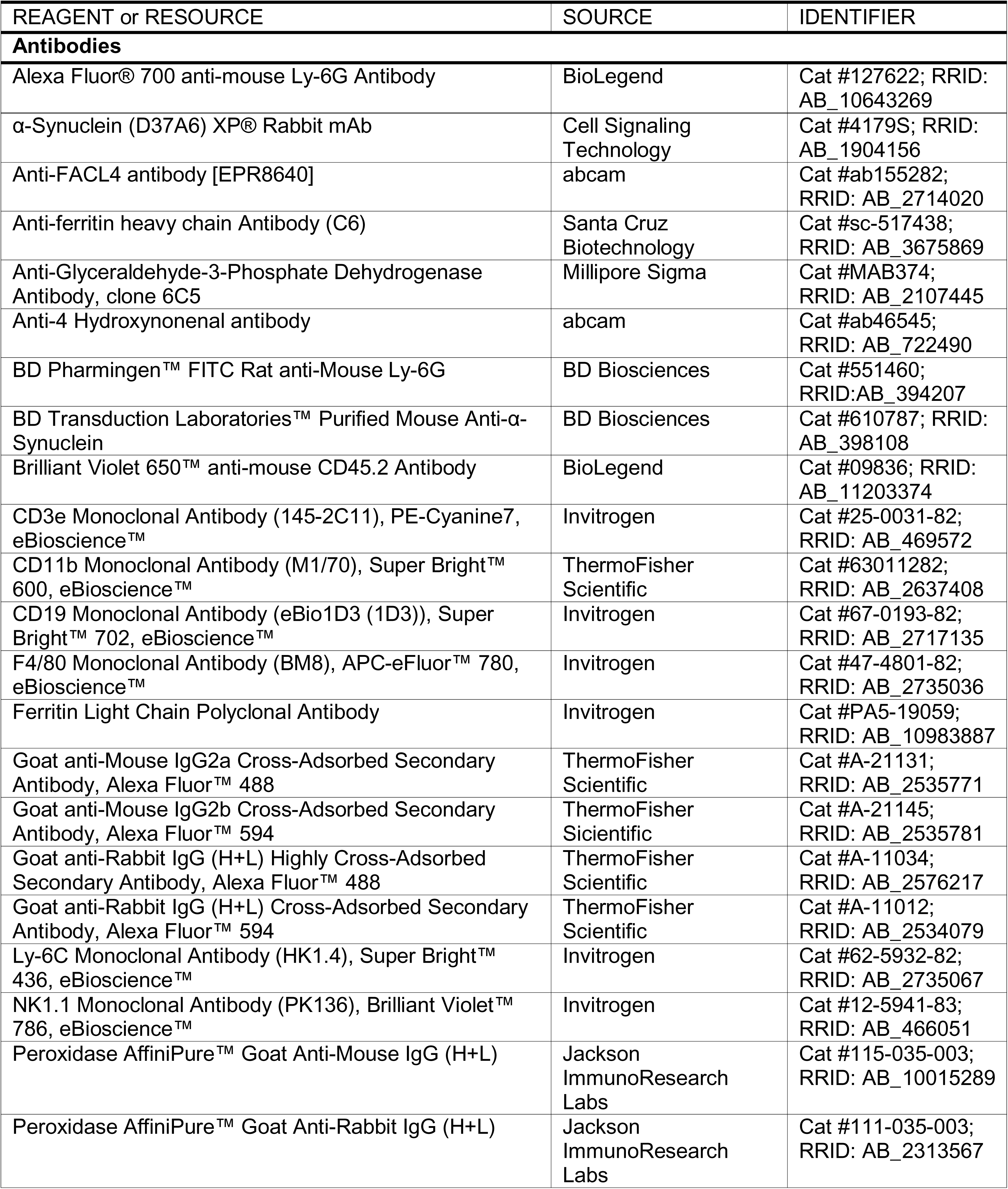
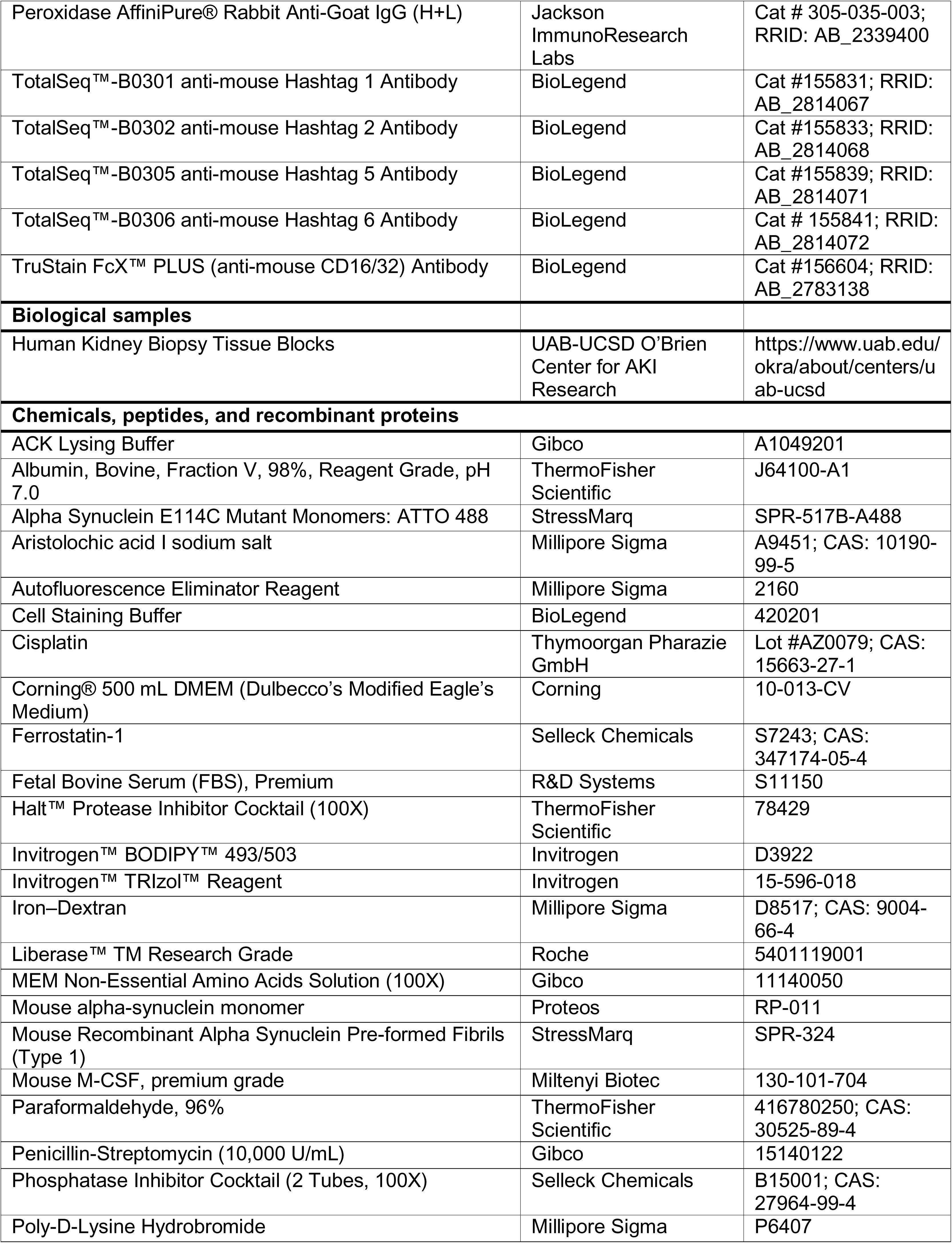
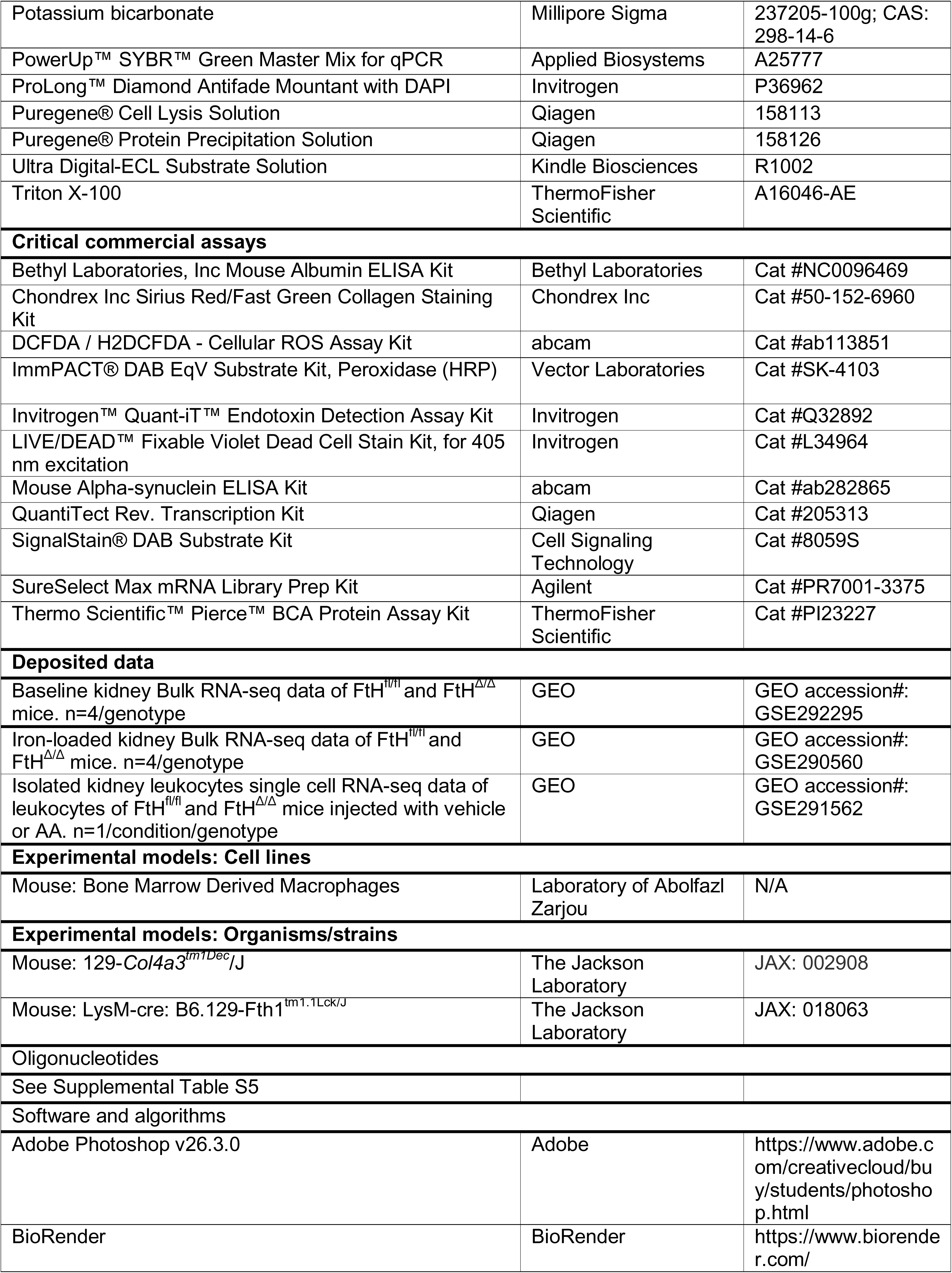
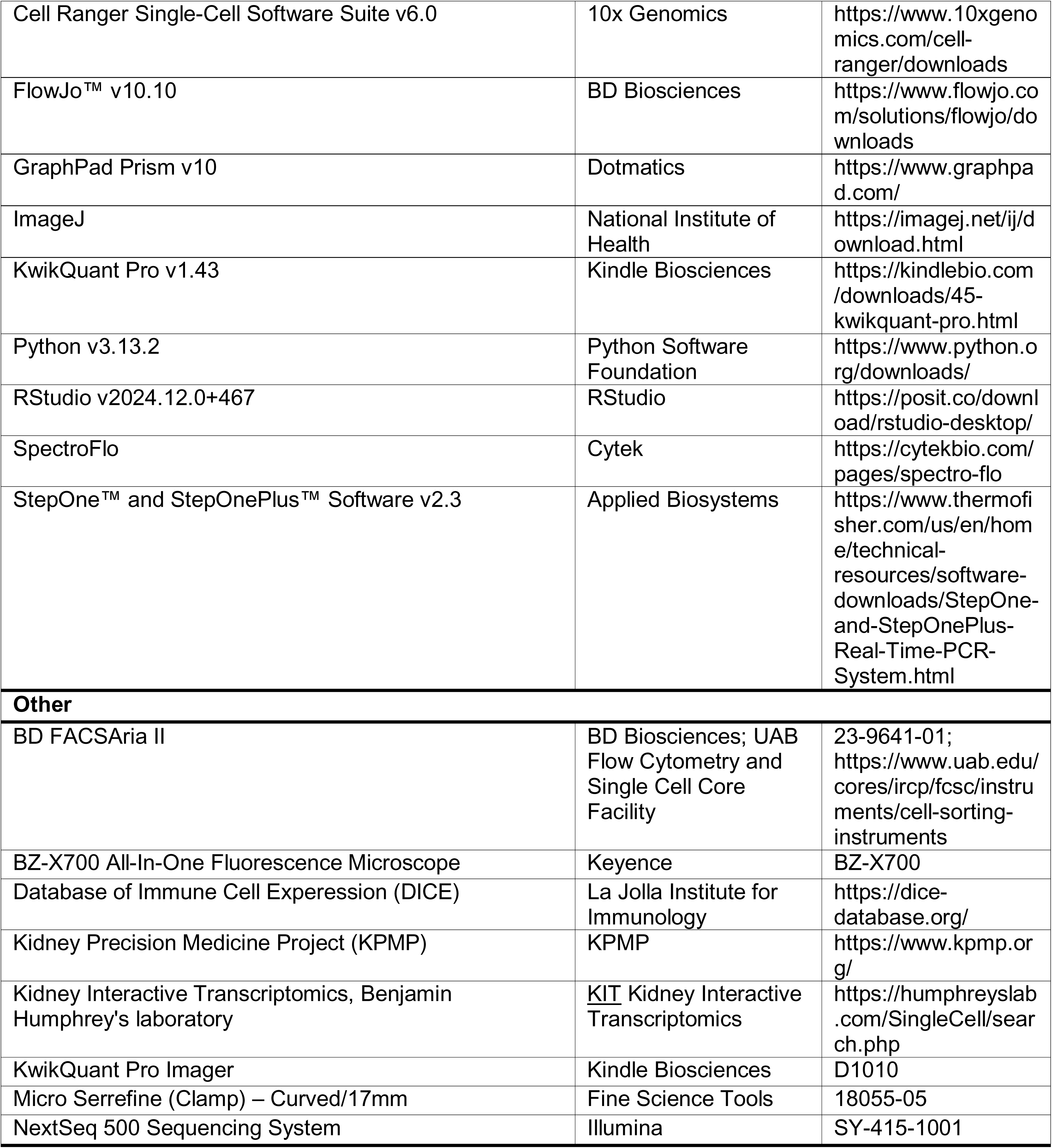

### Mice

Mice with targeted deletion of myeloid FtH were generated by crossing FtH-floxed mice (FtH^fl/fl^) mice^75^ with lysozyme 2-Cre mice that express the Cre protein in cells of myeloid lineage, generating littermates that are either FtH^fl/fl^ or FtH^Δ/Δ^ on a C57BL/6 background. Col4a3^+/+^ (wildtype) and Col4a3^−/−^ (Alport) from a mixed Sv129/C57BL/6 background were generated as previously described.^76^ Wildtype C57BL/6 mice were purchased from the Jackson Laboratory and the colony was expanded at UAB. All mice were maintained in temperature-controlled rooms in a 12-hour light/dark cycle and were provided with food and water ad libitum. Age of all experimental mice were between 8-12 weeks. Given the well-established resistance of female mice to kidney injury and progression of CKD, only male mice were used in this study. All animal experiments were approved by the Institutional Animal Care and Use Committee at UAB and were routinely monitored by the Animal Resource Staff.

### Aristolochic Acid (AA) model of kidney injury

Male FtH^fl/fl^ and FtH^Δ/Δ^ mice were randomly assigned to either the AA group or the control group. Mice in the AA group received intraperitoneal injections of AA (2 mg/kg body weight; Sigma-Aldrich, A9451) once daily for five consecutive days to induce kidney injury, while control mice received an equal volume of saline as a vehicle. Mice were harvested at weeks 1, 3, and 6 post-final injection, and organs, blood, and urine were collected for endpoint analysis.

### I/R model of AKI-to-CKD progression

Wild-type mice underwent a bilateral ischemia-reperfusion (I/R) renal injury (20 minutes ischemia time) procedure to induce AKI-CKD transition as described.^77^ Samples from a previous study were used in this report where temporal serum creatinine measurements were published.^77^ Briefly, mice were anesthetized with ketamine xylazine, and following incision, both renal pedicles were cross clamped for 20 minutes using an atraumatic vascular clamp (catalog #18055–05; Fine Science Tools, Foster City, CA). Immediate blanching of the kidney confirmed ischemic induction. Body temperature was maintained at 37°C during surgery and ischemia time. Kidneys were inspected for color change within 1 min of clamp removal to ensure uniform reperfusion. Animals were harvested at Day 1, 7, & 28 post-procedure.

### Cisplatin model of CKD

This model was established as previously described.^77^ Briefly, 9 mg/kg body weight cisplatin (1.0 mg/ml solution in sterile 0.9% saline) were administered intraperitoneally once a week for 4 weeks to induce moderate CKD and tissues were harvested 1 week post final injection. Control mice received saline as vehicle.

### Model of Alport syndrome

Col4a3^+/+^ (wild-type) and Col4a3^−/−^ (Alport) mice were harvested at 12 weeks of age, at which point substantial evidence of advanced CKD is present.^76^

### Parenterally induced systemic iron overload

Iron overload was established by injecting male FtH^fl/fl^ and FtH^Δ/Δ^ mice with 0.4mg/g bodyweight iron dextran (Millipore Sigma, D8517) once a day for 5 consecutive days as previously described.^78^ Three days after the final injection, mice were sacrificed for endpoint analysis.

### Ferrostatin-1 administration to mitigate ferroptosis

FtH^fl/fl^ and FtH^Δ/Δ^ mice were intraperitoneally injected with 1 mg/kg ferrostatin-1 (Fer-1, Selleck Chemicals, S7243) on day −1, one day before first AA injection. Following first AA administration, mice continued to receive Fer-1 injections three times a week.

### Kidney function measurement

Serum and urine creatinine were quantified at the UAB-UCSD O’Brien Center for AKI research using liquid chromatography–tandem mass spectrometry (LC-MS/MS) as previously described.^79^

### Western blot analysis

Tissues and cells were homogenized, and protein was isolated as previously described.^79^ Lysates were centrifuged at 15,000 g for 15 min, and the supernatant was collected. Protein concentrations were measured by BCA assay kit (ThermoFisher Scientific, PI23227). Protein samples were denatured at 95°C for 5 minutes for standard westerns, or at 50°C for 30 minutes for Snca westerns, and then loaded on a 12% Tris-glycine sodium dodecyl sulfate polyacrylamide electrophoresis gel. Western gels were then transferred onto 0.45 um PVDF transfer membranes (ThermoFisher Scientific, 88518) via electroblotting. After transfer, Snca westerns were incubated for 30 minutes in 0.4% paraformaldehyde in 1X PBS before blocking for 1 hour in 6% non-fat dry milk in PBS with 0.1% Tween-20 (PBST). All other blots were immediately blocked in 6% non-fat dry milk in PBST for one hour after transfer. After blocking, membranes were then incubated overnight at 4°C in 1% non-fat dry milk in PBST with primary antibody (dilutions for primary antibodies can be found in Table S4). Secondary antibodies of anti-Rabbit or anti-mouse HRP-conjugated (JacksonImmuno Research Labs, 115-035-003 & 111-035-003) were applied at 1:2500 dilutions in 5% non-fat dry milk in PBST for one hour at room temperature. HRP-conjugated antibodies were detected via chemiluminescence using Ultra Digital-ECL Substrate Solution (Kindle Biosciences, R1002) and developed with the KwikQuant Pro Imager (Kindle Biosciences, D1010). Specificity of monomeric Snca bands was confirmed in preliminary studies using brain tissue and recombinant Snca as positive controls. Densitometry analysis was performed using Image J 1.54k (NIH) and results were normalized to GAPDH. Additional Western blots used for densitometry represent independent samples and are provided, along with representative blots, as a supplemental file.

### Quantitative realtime-PCR (qRT-PCR) to measure mRNA expression

Total RNA was extracted from cells and tissues using TRIzol (Invitrogen,15596018). cDNA was synthesized from total RNA with the QuantiTect Rev. transcription kit (Qiagen, 205313) per the manufacturer’s instructions. Quantitative-PCR was performed using PowerUp™ SYBR™ Green Master Mix (Applied Biosystems, A25777) on Applied Biosystems StepOnePlus™ Real-Time PCR System. Quantification of gene expressions was done via the 2-^ΔΔ^Ct method, and GAPDH expression was used for normalization. All real-time primers can be found in Table S5. All reactions were performed in triplicate, and specificity was monitored using melting curve analysis.

### Immunohistochemistry (IHC) staining

IHC staining was performed as previously described.^80^ Kidneys were fixed in 10% neutral buffered formalin for 24 h and then embedded in paraffin. Paraffin-embedded 5 μm kidney sections were deparaffinized in xylenes, rehydrated in a series of ethanol rinses from 100-70% ethanol, then washed in distilled water. Antigen retrieval was performed in Trilogy (Cell Marque, 922P-06) at 95°C for 30 min. Sections were allowed to cool slowly, washed in distilled water, and incubated in 3% H_2_O_2_ for 20 min. Sections were blocked in blocking buffer containing 1% BSA (ThermoFisher Scientific, J64100-A1), 0.2% non-fat dry milk, and 0.3% Triton X-100 (ThermoFisher Scientific, A16046-AE) in 1X PBS for 30 minutes. The primary antibody, Snca (Cell Signaling, 4179S), was diluted 1:200 in the blocking buffer and incubated overnight at 4°C. Sections were washed once with PBST and twice with PBS for 5 minutes each. Anti-rabbit HRP-conjugated secondary antibody (Jackson ImmunoResearch Labs, 115-035-003) was diluted 1:500 in blocking buffer and added to the sections for 1 hour at room temperature. Sections were washed once with PBST and twice with PBS for 5 min each. Chromogen substrates (Vector Labs, SK-4103) were mixed per the manufacturer’s instructions and added to sections. Sections were washed in distilled water, dehydrated, and mounted using xylene mounting media. Images were captured on a BZ-X700 All-In-One Fluorescence Microscope (Keyence, Istasca, IL).

### Immunofluorescence staining

Deparaffinization, rehydration, antigen retrieval, blocking was similar to IHC protocol outlined above. Sections were then incubated overnight at 4°C with primary antibodies (primary antibodies dilutions can be found in Table S4) diluted in blocking buffer. Sections were washed once with PBST and twice with PBS for 5 minutes each and then incubated with fluorescently tagged secondary antibodies (see Key Resource Table) at 1:500 dilutions in blocking buffer. Sections were washed once with PBST and twice with PBS for 5 minutes each, followed by application of autofluorescence eliminator reagent (Millipore Sigma, 2160) per manufacturer’s instructions. Slides were then mounted with DAPI (Invitrogen, P36962) and stored at −30°C until they were ready for viewing. Images were captured on a BZ-X700 All-In-One Fluorescence Microscope (Keyence, Istasca, IL). Notably, the very intense staining of Snca in MΦ required us to decrease the intensity of staining to better depict these MΦ at expense of tubules. Brightness, and contrast adjustments were made in an identical manner for matched sections and was applied to the entire image.

### Sirius red/fast green stain

Mouse kidneys were fixed in 10% neutral buffered formalin for 24 h and then embedded in paraffin. Sirius Red/Fast Green Stain was performed on the tissue sections using a sirius red/fast green collagen staining kit (Chondrex, 50-152-6960) per the manufacturer’s instructions, then dehydrated and mounted with xylene mounting medium. Four images per kidney were acquired at a 10x magnification with a BZ-X700 All-in-One Fluorescence Microscope. Percent area positive and intensity (integrated density) of Sirius red stain, as a measure of collagen deposition and fibrosis, was measured using ImageJ using the “Triangle” auto-threshold method. Color, brightness, and contrast adjustments were made in an identical manner for matched sections.

### Iron deposition staining

Kidneys were fixed in 10% neutral buffered formalin for 24 h and then embedded in paraffin. Paraffin-embedded 5 μm kidney sections were deparaffinized in xylenes, rehydrated as mentioned above. Sections were then placed in a working solution of distilled water containing 4% HCl and 4% potassium ferrocyanide for 30 minutes at room temperature, rinsed in tap water, and then dehydrated and mounted with xylene mounting media to achieve a Perl’s Prussian blue stain. For enhanced iron deposition staining, the modified Perl’s stain method was implemented. After rehydration, sections were blocked in 1% H_2_O_2_ for 10 minutes and then washed with distilled water. Slides were then incubated for 1 hour at room temperature in a working solution of distilled water containing 4% HCl and 4% potassium ferrocyanide. Sections were then washed with distilled water, and then chromogen substrate (Cell Signaling, 8059S) was applied. Sections were washed in distilled water, dehydrated, and mounted using xylene mounting media.

### Flow cytometry and fluorescence-activated cell sorting (FACS)

Leukocytes (CD45^+^ cells) were isolated from kidneys as previously described.^81^ Mice were anesthetized with isoflurane and perfused through the left ventricle with 10 mL of cold PBS. Kidneys were then removed, stripped of the capsule, weighed, and minced with a razor blade on a glass slide. The minced tissue was placed in 2 mL of DMEM containing 1.67 Wunsch units/mL Liberase (Roche, 5401119001) and incubated at 37°C for 30 minutes. Digestion was halted by adding cold PBS containing 1% BSA. The resulting suspensions were passed through an 18-gauge needle three times and subsequently filtered through a 40-µm strainer. The samples were then centrifuged at 400g for 7 minutes, and the supernatant was aspirated. Red blood cells were lysed using ACK lysis buffer for 2 minutes at room temperature, followed by washing with ice-cold PBS to obtain the remaining leukocytes. Cells were then stained with a violet fixable viability dye (Invitrogen, L34964) for 20 minutes in dark at room temperature and treated with an unlabeled anti-CD16/32 antibody (BioLegend, 156604) to block Fcγ3 receptors for 30 minutes. Subsequently, cells were stained with the following antibodies: anti–CD45.2 Brilliant Violet 650 (BioLegend, 09836), anti-CD11b Super Bright 600 (M1/70, Invitrogen), anti-F4/80 APC-eFluor-780 (Invitrogen, 63011282), anti-NK1.1 PE-C7 (Invitrogen, 12-5941-83), anti-Ly6G Alexa Fluor 700 (BioLegend, 127622), anti-Ly6C Super Bright 436 (Invitrogen, 62-5932-82), anti-CD19 Super Bright 702 (BioLegend, 67-0193-82), and anti-CD3e PE-Cy7 (Invitrogen, 25-0031-82). Following staining, samples were washed and resuspended in 500 µL of staining buffer before analysis by flow cytometry using the Cytek Northern Lights instrument and SpectroFlo software. Single-color controls were used for spectral unmixing, and fluorescence-minus-one (FMO) controls were applied to set positive gates. Data were analyzed using FlowJo software.

### Bulk RNA sequencing data processing and analysis

Total RNA was isolated from kidneys using TRIzol. RNA quality was assessed using the Agilent 2100 Bioanalyzer and only samples with a RIN score > 8.0 were used for RNAseq. RNA was sequenced on NextSeq500 system (Illumina), and the library was prepared with the Agilent SureSelect Stranded mRNA kit. STAR (version 2.7.7a) was used to align the raw RNA-Seq fastq reads to the mouse reference genome (GRCm38 p6, Release M25) from Gencode.^82^ Following alignment, HTSeq-count (version 0.11.3) was used to count the number of reads mapping to each gene.^83^ Normalization and differential expression were then applied to the count files using DESeq2.^84^ Heatmaps were generated using publicly available Morpheus platform. (https://software.broadinstitute.org/morpheus/)

### Single-cell RNA sequencing

CD45⁺ immune cells were isolated by FACS from the kidneys of vehicle- and AA-treated mice at 6 weeks post final injection, following the flow cytometry protocol described earlier. CD45^+^ cells were sorted on a BD FACSAria II at the UAB Flow Cytometry and Single Cell Core Facility. Isolated cells were then processed using the Chromium 3′ Single Cell RNA sequencing kit (10× Genomics) in accordance with the manufacturer’s protocol. Cell-specific bar-coded sequencing libraries were generated, and sequencing was performed on an Illumina NovaSeq6000. Reads were processed using the 10× Genomics Cell Ranger Single-Cell Software Suite (v.6.0). Each sample of scRNA-seq was mapped to the mouse reference genome (GRCm38) provided by 10x Genomics. The samples then were merged by loading the Cell Ranger output h5 file using Seurat (v 5.1) for data preprocessing, integration, and normalization. Cells with fewer than 200 genes, more than 6,000 unique molecular identifiers (UMIs), and > 10% mitochondrial UMIs were excluded. RNA expression was normalized by a scaling factor of 10,000 and log-transformed (base 2), and the top 2500 variable genes were scaled via the ScaleData function. Normalization returned two gene-cell matrices: one in log scale, and the other the adjusted gene-cell count. PCA analysis was performed with 30 principal components (PCs) via the RunPCA function, followed by Harmony analysis to normalize the PCA embeddings across samples. We also used 15 as the number of dimensions used for the FindNeighbors function. We ran the FindClusters function (resolution = 0.6) to identify clusters of cells and the RunUMAP function (reduction = “harmony”, dims = 1:20) for reduction to 2 dimensions for visualization purposes. The assignment of each cluster to a specific cell type was done with SingleR package using the ImmGenData reference dataset. Differentially expressed genes between two groups were identified using the Findmarkers function with the default Wilcoxon’s rank-sum test and genes with adjusted p-value <0.05 were considered differentially expressed. Pathway analysis of differentially expressed genes: differentially expressed genes for each cell-type were functionally annotated using the R package clusterProfiler (version 3.14.0). The mouse Kyoto Encyclopedia of Genes and Genomes (KEGG) and gene ontology (GO) databases were used to determine associations with particular biological processes, diseases, and molecular functions. The top pathways with an FDR-adjusted p-value <5% were summarized in the results.

### Measurement of serum and urinary Snca

Serum Snca levels were measured using the Mouse Alpha-synuclein ELISA Kit (Abcam, ab282865) per the manufacturer’s protocol. Based on preliminary results serum from FtH^fl/fl^ mice was diluted 10x for both vehicle and CKD cohorts, while FtH^Δ/Δ^ serum was diluted 300x for both vehicle and CKD cohorts. Urine from vehicle treated mice were diluted 5x, while urine from CKD mice were diluted 20x. Urinary Snca levels were normalized to creatinine.

### Quantification of albuminuria

Urinary albumin was quantified using the Mouse Albumin ELISA kit (Bethyl Laboratories, NC0096469), according to the manufacturer’s protocol. Data were normalized to urine creatinine.

### Cell culture

Bone marrow derived macrophages (BMDMs) were isolated from wild-type mice as previously described.^85^ BMDMs were cultured in 6-well plates (Corning, 3471) for 5 days in DMEM (Corning, 10-013-CV) with 15% FBS (R&D Systems, S11150), 1% penicillin/streptomycin (Gibco, 15140122), 1% MEM non-essential amino acids (Gibco, 11140050), and 0.3 µg/ml M-CSF (Miltenyi Biotec, 130-101-704). On Day 6, the culture media was replaced with DMEM containing 1% FBS and 1% penicillin/streptomycin for 2 hours, followed by treatment with either vehicle, mouse recombinant monomeric Snca protein monomers (Proteos, RP-011), or fibrils at a dose of 1µM (Stressmarq, SPR-324). Dose of Snca was established per previous reports. Cells were then washed with PBS and collected for RNA or protein analysis. Presence of LPS mouse recombinant Snca monomers and fibrils was ruled out using the Pierce chromogenic endotoxin quantification kit (ThermoFisher, A39552) according to the manufacturer’s instructions where values were determined to be < 0.01 EU/ml. Fibrils were sonicated at 4°C in a water bath sonicator for a total of 15 minutes, using a cycle of three seconds on and two seconds off. To assess the effects of heat and enzymatic digestion on Snca, 1µM of monomers were either boiled overnight at 100°C or treated with proteinase-K (1 mg/mL) at 37°C overnight before addition to BMDMs media.

### Live cell imaging for Snca uptake

For the uptake assay, BMDMs plated on 35 mm MatTek live imaging dishes and were treated with 0.1µM of Alexa 488 (A488)-labeled Snca monomers (StressMarq, SPR-517B-A488). At 0, 20, 30, 60, and 120 minutes following A488-labeled Snca treatment, extracellularly bound Snca was quenched using trypan blue. Both A488 and Alexa 555 channels were used for imaging. A488 was used to visualize Snca internalization and Alexa 555 to detect trypan blue binding to extracellular proteins. The internalization of A488-labeled Snca was then assessed via live cell imaging, as previously described.^86^

### BODIPY (lipid peroxidation sensor) assay

BMDMs were treated with either vehicle or monomeric Snca at a concentration of 0.1 or 1 µM) (Proteos, RP-011) for 16 hours and incubated with 2 µM BODIPY™ 581/591 C11 (Invitrogen, D3922) for 30 minutes at 37°C in 5% CO_2_. Following incubation, cells were washed with cold PBS to remove excess dye and resuspended in flow cytometry buffer (PBS containing 2% FBS and 0.1% sodium azide). Flow cytometry was performed using the Cytek Northern Lights instrument and analyzed using FlowJo software.

### Cellular ROS assay

BMDMs were treated with either vehicle or Snca (0.1 or 1 µM) monomers (Proteos, RP-011) for 16 hours and incubated with 10 µM DCFDA (Abcam, ab113851) for 30 minutes at 37°C in 5% CO₂. After incubation, cells were washed with cold PBS to remove excess dye, followed by fluorescent microscopy and flow cytometry analysis. Flow cytometry was performed using the Cytek Northern Lights instrument and analyzed using FlowJo software.

### Metallomics quantification: Inductively coupled plasma mass spectrometry (ICP-MS)

ICP-MS measurements were performed by the Iron and Heme Core Facility at the University of Utah (grant number: U54DK110858) to measure tissue iron, copper, zinc, and magnesium content. Water to about 3 to 4 times the weight of tissue was added and then sonicated using a Fisher FB505 Sonic Dismembrator with a 1/16” microtip. A Pierce BCA Protein Assay Kit (Thermo Scientific) was used to determine protein concentration in the resulting homogenates. Triplicate 25 µL aliquots per sample were digested in open acid-washed tubes overnight with 500 µL of optima grade 70% nitric acid (Fisher Scientific) and 100 µL ultrapure 30% hydrogen peroxide (Fisher Scientific). Background blanks contained just water. The tubes were covered with big glass dish to protect from dust during digestion. The mixtures were then dried on a heat block at 98°C, cooled and dissolved in 2% HNO3 containing 100 ppb Ge internal standard. The solutions were next put into 96-position deep-well plates and analyzed for 56Fe, 24Mg, 63Cu, 66Zn and using an Agilent 7900 ICP-MS system, with helium as collision gas. The Agilent Multi-Element Calibration Standard 2A was diluted to various concentrations from 0.005ppb to 2700ppb in 3-fold dilution steps to serve as reference for quantification. An Agilent SPS4 autosampler was used to deliver the samples from the plate to the ICP-MS spectrometer. This system was also equipped with a MicroMist glass concentric nebulizer, quartz spray chamber (cooled to 2°C), quartz torch with 2.5 mm internal diameter injector and standard Nickel interface cones. Agilent ICP-MS MassHunter 4.4 software was used to control the instrument and process raw data. Some run parameters were as follows: RF Power at 1600 W, carrier gas at 1.05 L/min, sample flow rate at 0.2 mL/min and collision cell gas flow rate at 5 mL/min. All results were normalized to protein content.

### Human samples

Remnant human kidney biopsies were obtained through the tissue resource available from the UAB-UCSD O’Brien Center for AKI Research. These samples were collected, de-identified, archived together, and analyzed according to protocols approved by the Institutional Review Board of the UAB. Biopsy samples for this study were selected based on the diagnosis of ACR, AIN, or TBM, according to pathologists’ diagnoses and comments. As controls, we used kidney biopsies from patients with TBM disease. Diagnosis of TBM was also confirmed by electron microscopy, and thickness measurement values used to establish the diagnosis of TBM were as follows: 337 ± 75 nm for females and 355 ± 75 nm for males (Figure S11). The three studied groups, TBM, AIN, ACR were each represented by kidney biopsies from six patients. Patient characteristics are presented in Table S1. Serum creatinine of one patient in TBM, and one patient in AIN group was not available for analysis.

### Interpretation of gene expression data using publicly available transcriptomic platforms

Data presented in this study received written permission from all resources mentioned below. We used three publicly available resources for transcriptomic analysis. These include the following. 1) Kidney Precision Medicine Project (https://www.kpmp.org). Accessed between September 2024 and February 2025. Funded by the National Institute of Diabetes and Digestive and Kidney Diseases (grant numbers: U01DK133081, U01DK133091, U01DK133092, U01DK133093, U01DK133095,U01DK133097, U01DK114866, U01DK114908, U01DK133090, U01DK133113, U01DK133766, U01DK133768, U01DK114907, U01DK114920, U01DK114923, U01DK114933, U24DK114886, UH3DK114926, UH3DK114861, UH3DK114915, UH3DK114937). Specifically, Figure S9 and Table S1, 2 were generated via accessing this platform. 2) Kidney interactive transcriptomics platform (http://humphreyslab.com/SingleCell/), developed by Dr. Humphreys. The generated results in this study used a data source that was previously published.^87,88^ Figure 7K, L and Figure S10 were generated using this platform. 3) Dice data base website (https://dice-database.org) was utilized to generate data presented in Figure S3.

### Statistical analysis

Experiments involving more than two groups were analyzed by 1-way or 2-way ANOVA, followed by post-hoc Dunnett’s, Sidak’s, or Tukey’s multiple comparison test. Statistical analyses were performed using appropriate methods based on the study design. Experiments involving two groups were analyzed by two-tailed, unpaired student’s t-test. For analyses involving three or more independent groups, a one-way ANOVA with Tukey’s or Dunnett’s post hoc correction was applied. When comparing two groups across two conditions, a two-way ANOVA with Tukey’s or Sidak’s correction for multiple comparisons was conducted. All data are expressed as mean ± SEM. Statistical tests were two-sided and performed using GraphPad Prism v10, with details provided in the figure legends. Statistical significance was defined as follows: ns (not significant, p > 0.05); *p < 0.05; **p < 0.01; ***p < 0.001.

