## Supplemental Figures for "Myeloid FtH Regulates Macrophage Response to Kidney Injury by Modulating Snca and Ferroptosis"

### Supplemental Figure S1

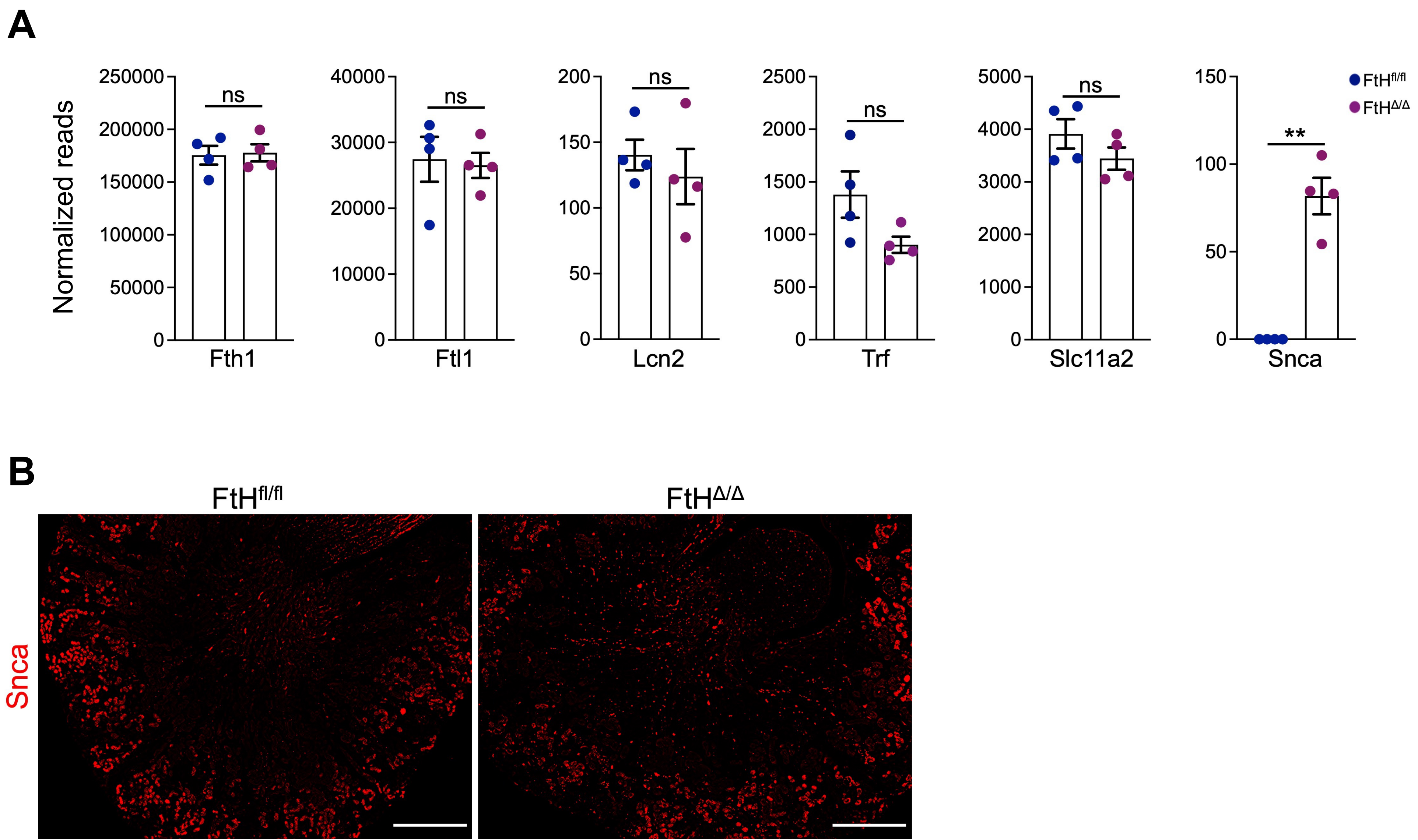

### Supplemental Figure S2

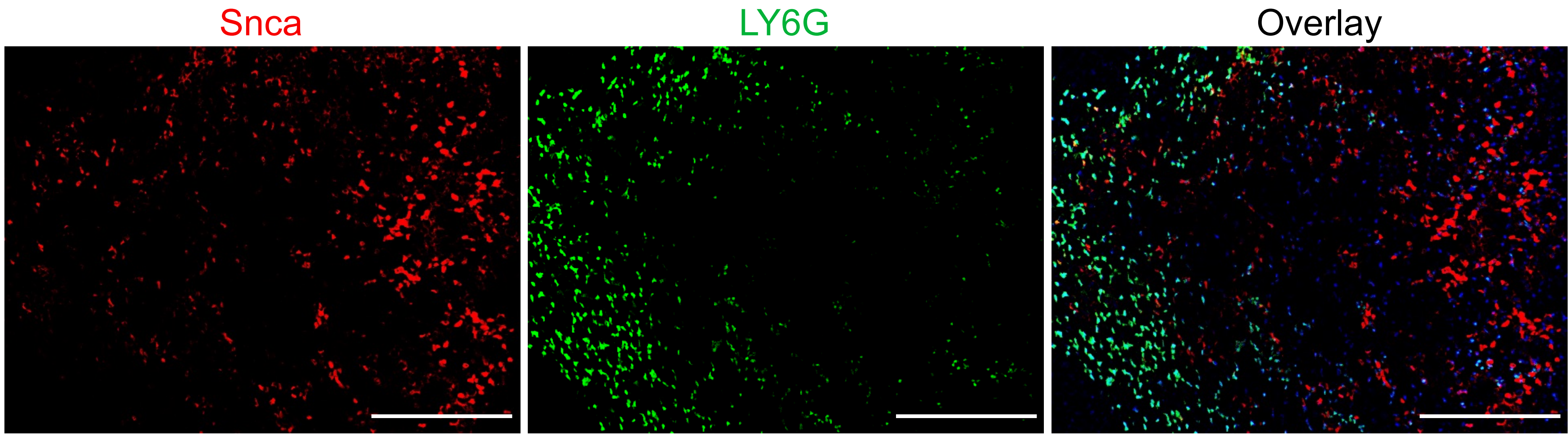

### Supplemental Figure S3

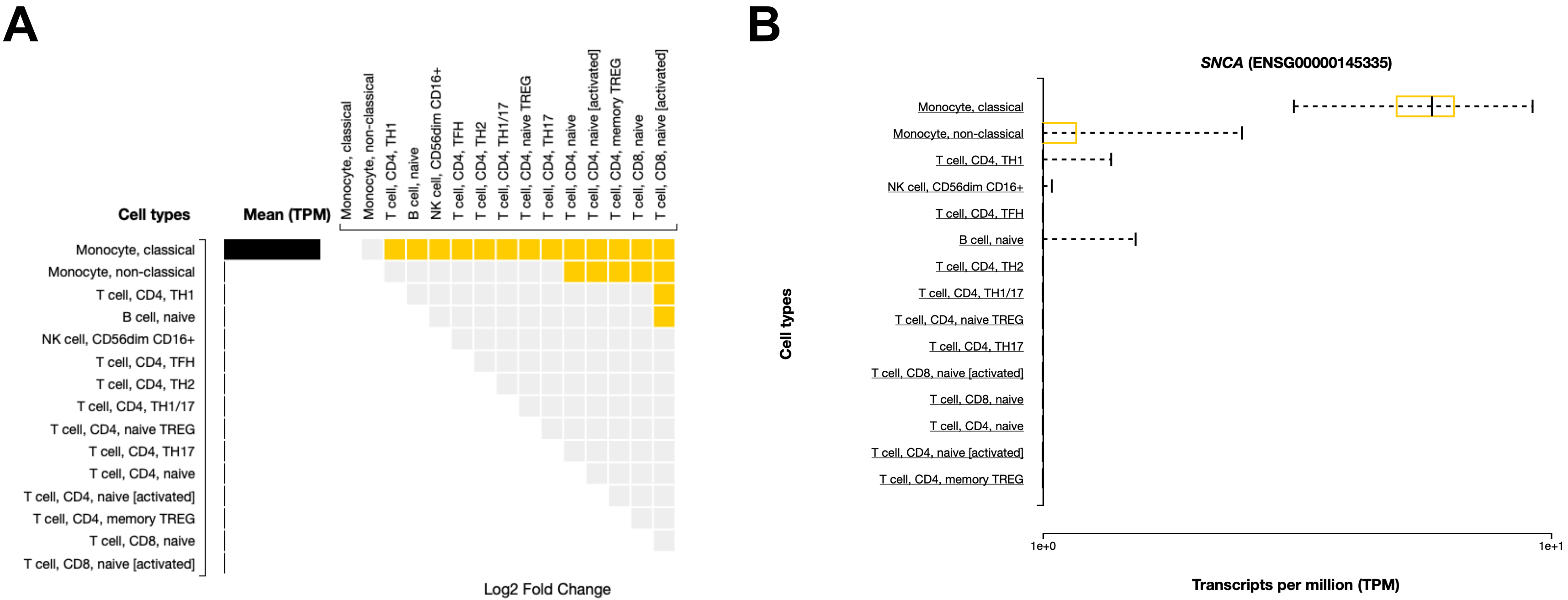

### Supplemental Figure S4

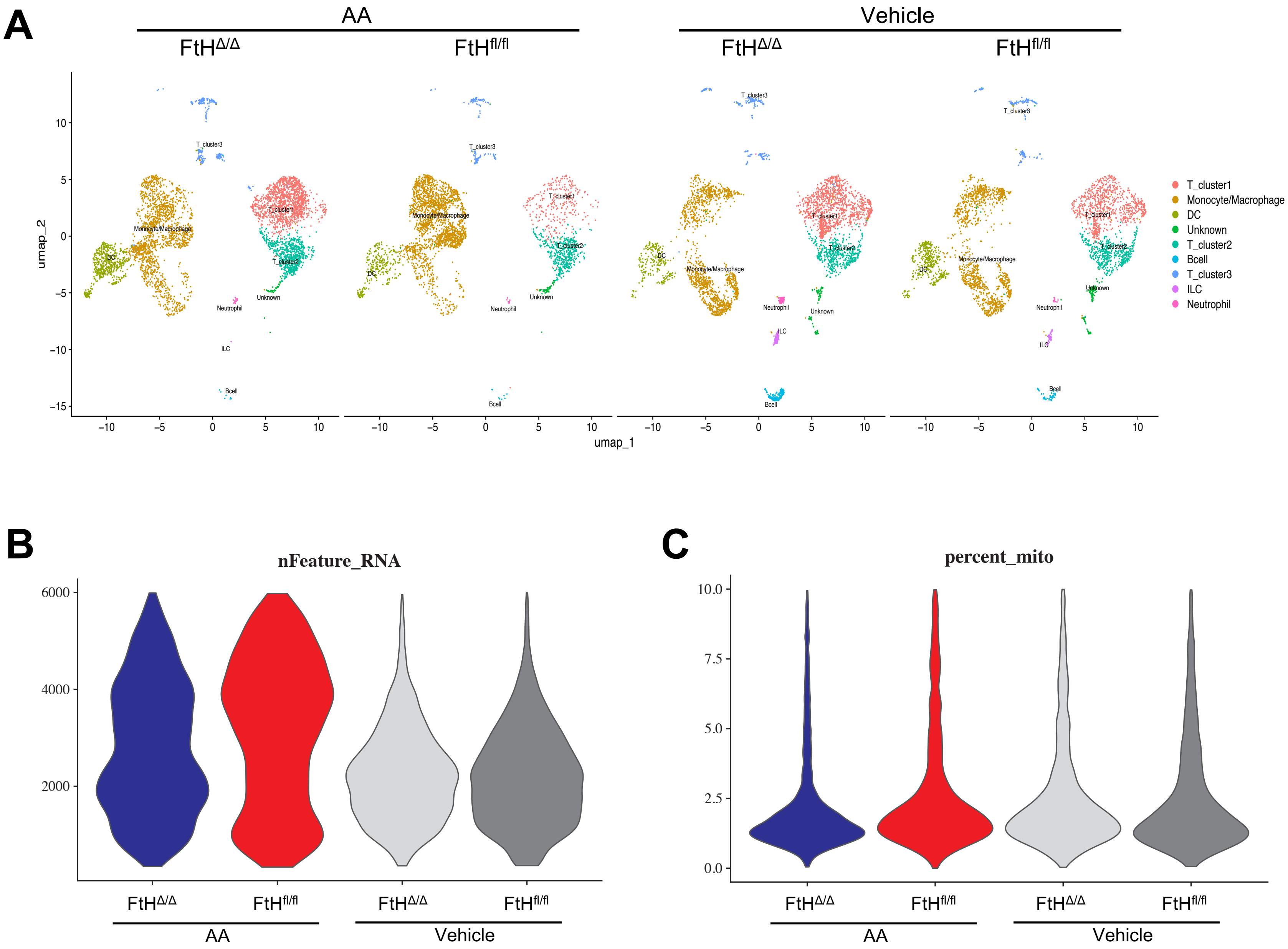

### Supplemental Figure S5

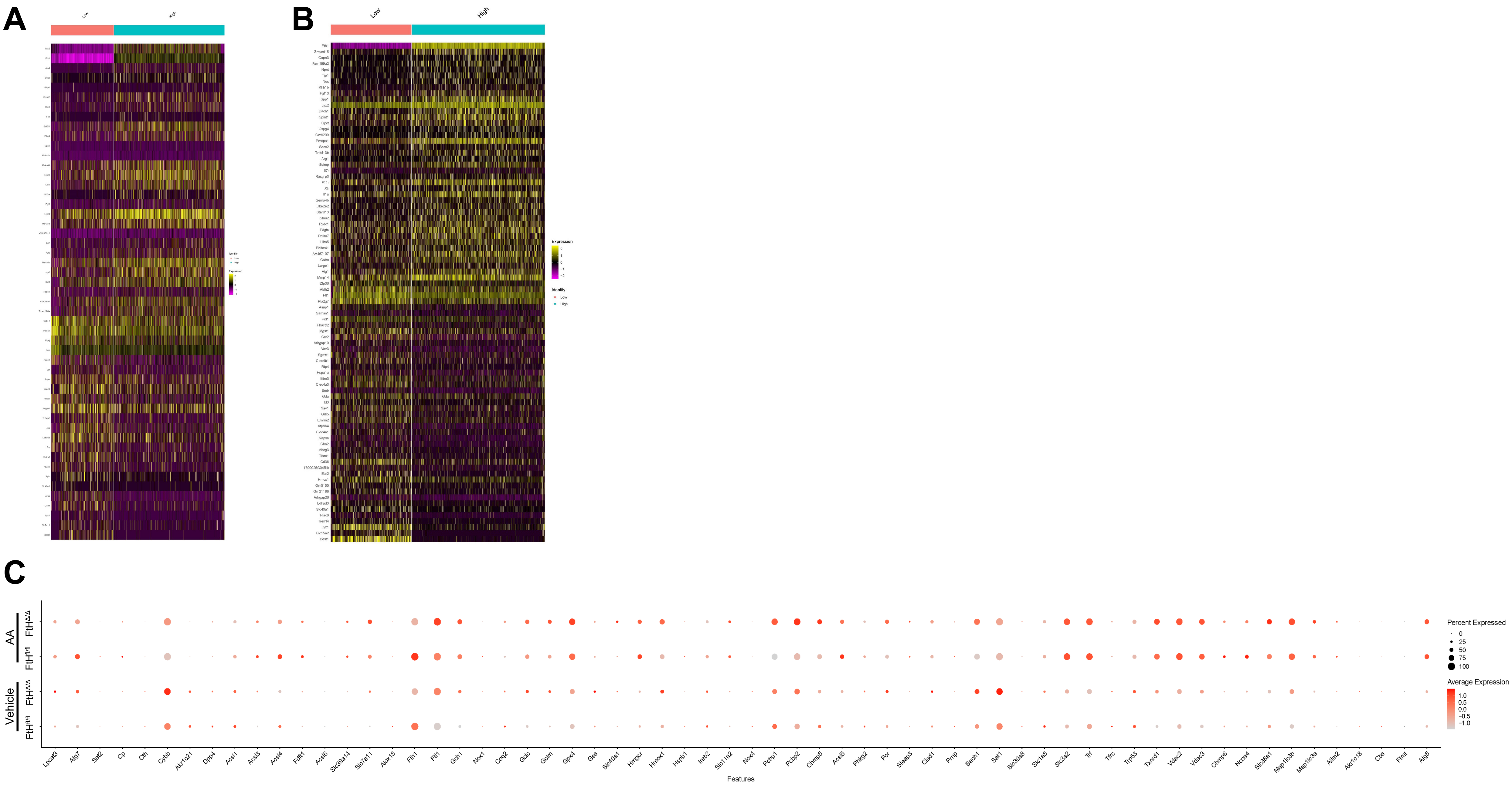

### Supplemental Figure S6

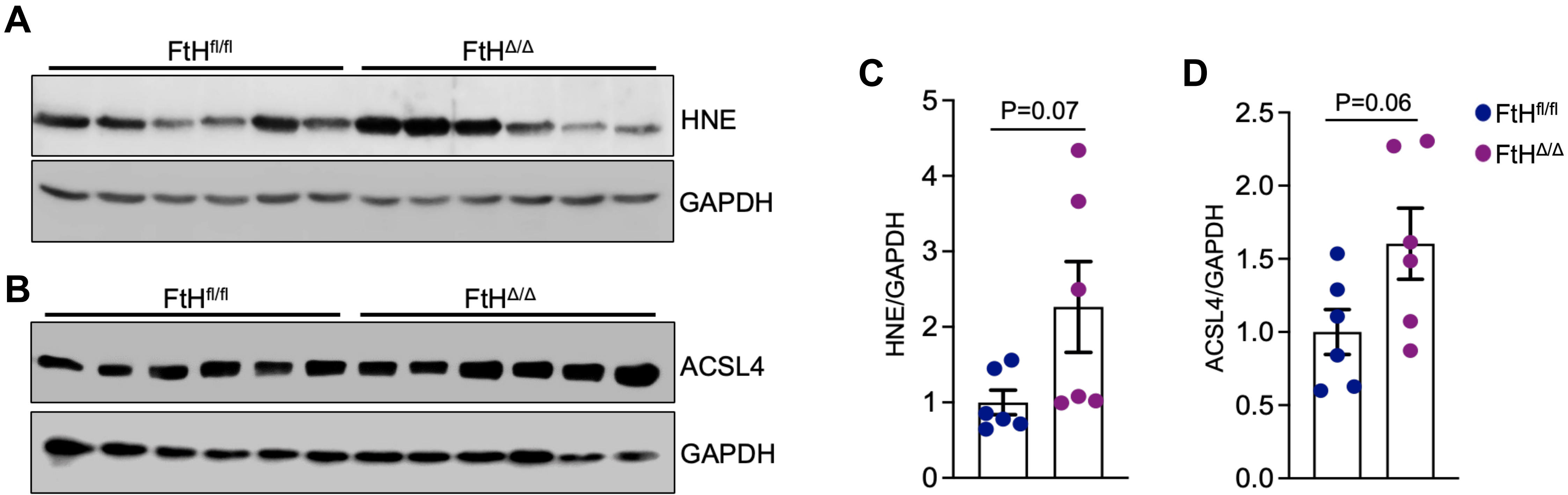

### Supplemental Figure S7

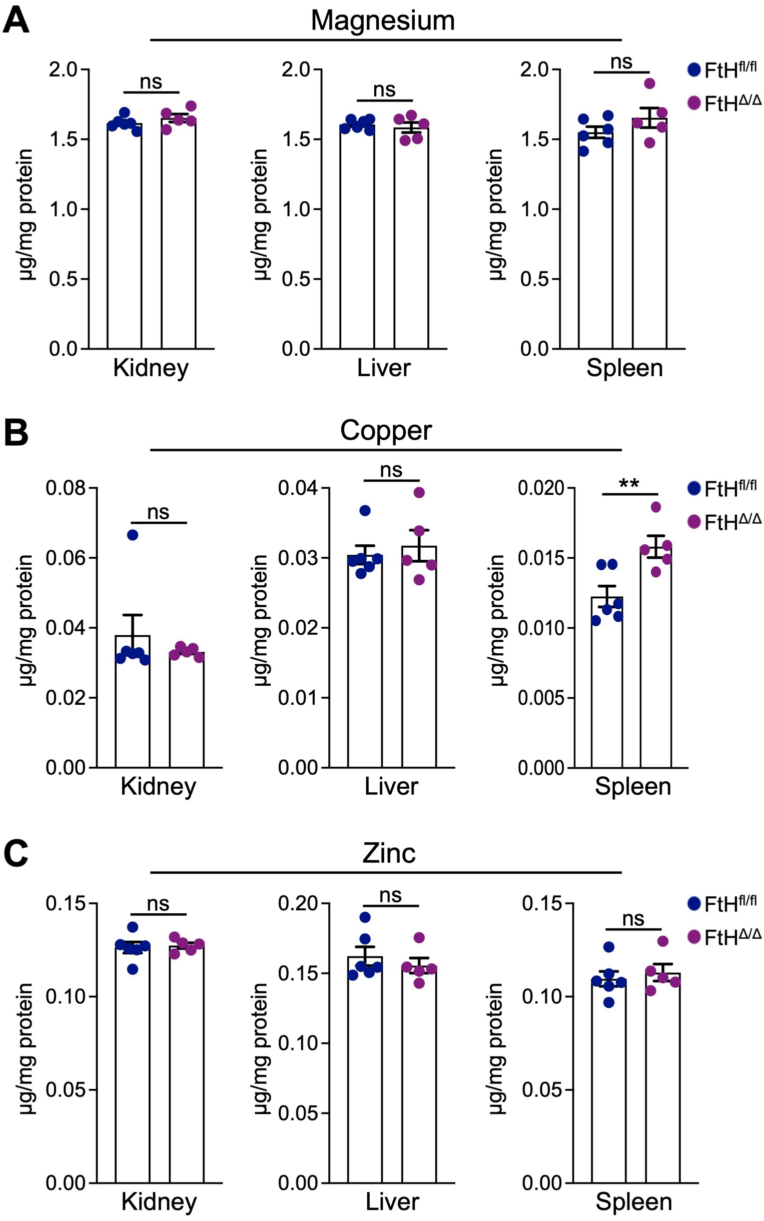

### Supplemental Figure S8

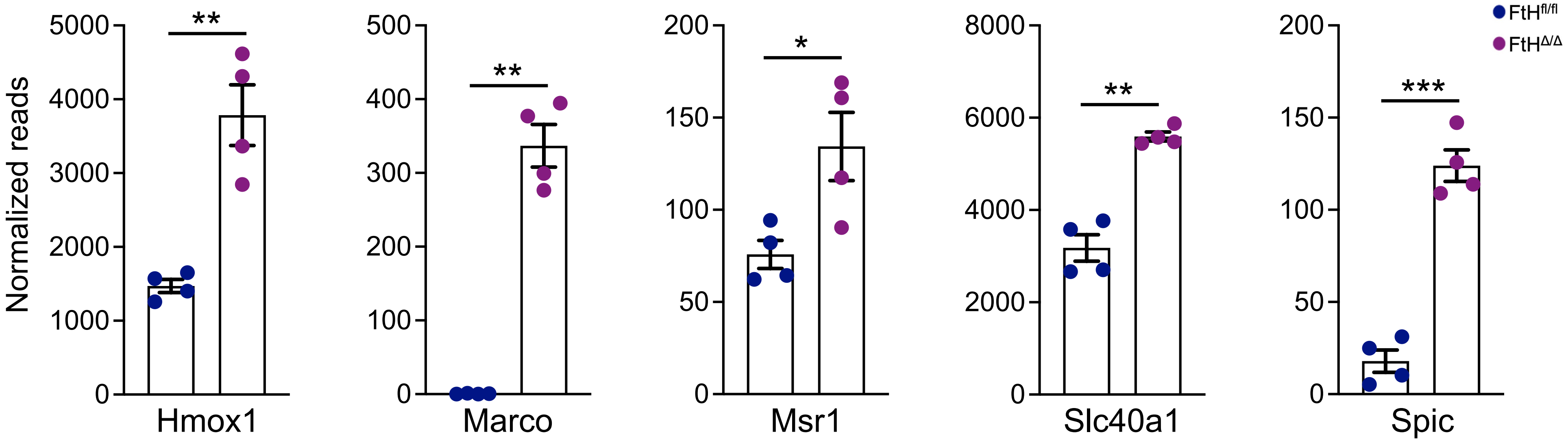

### Supplemental Figure S9

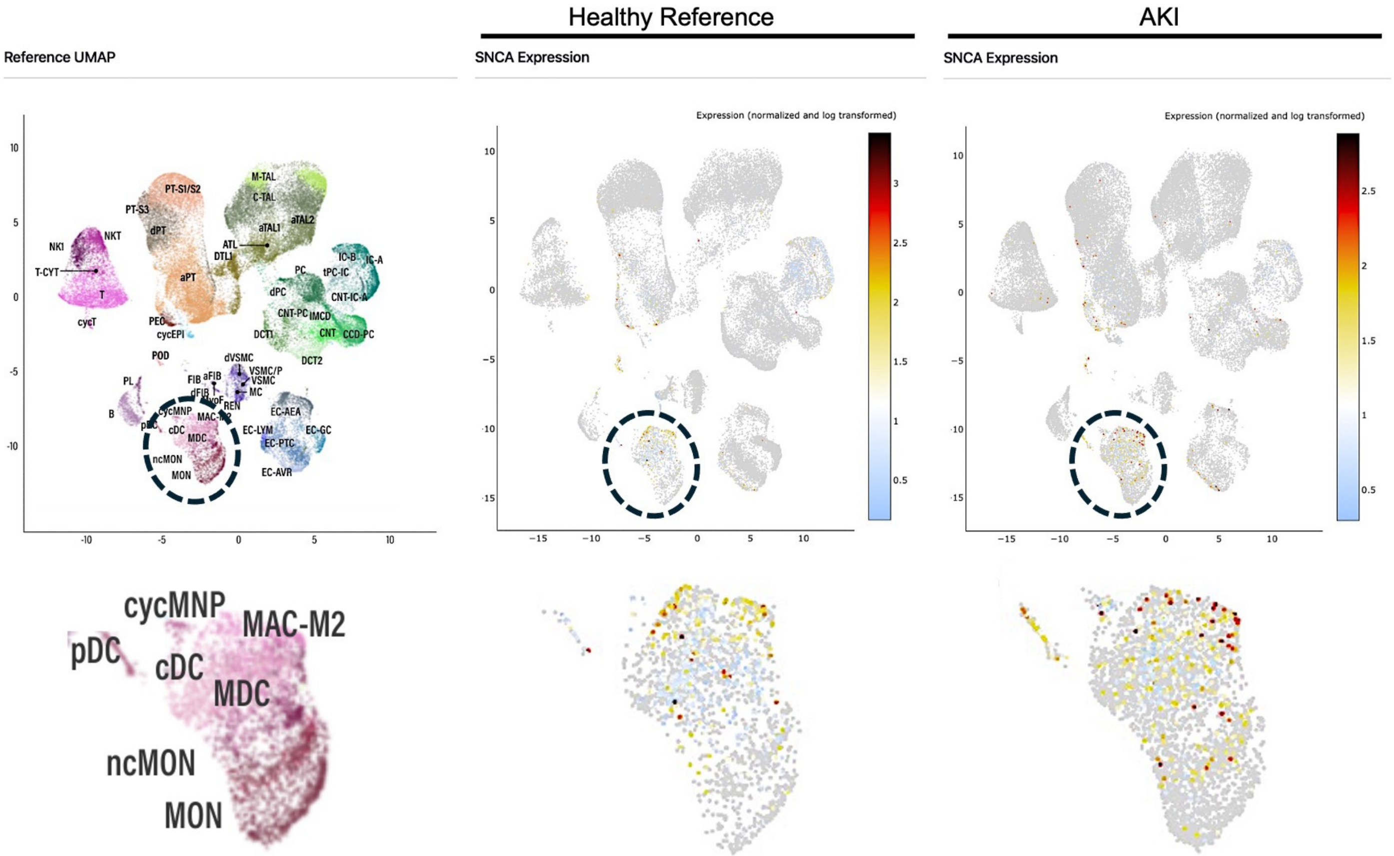

#### Supplemental Figure S10

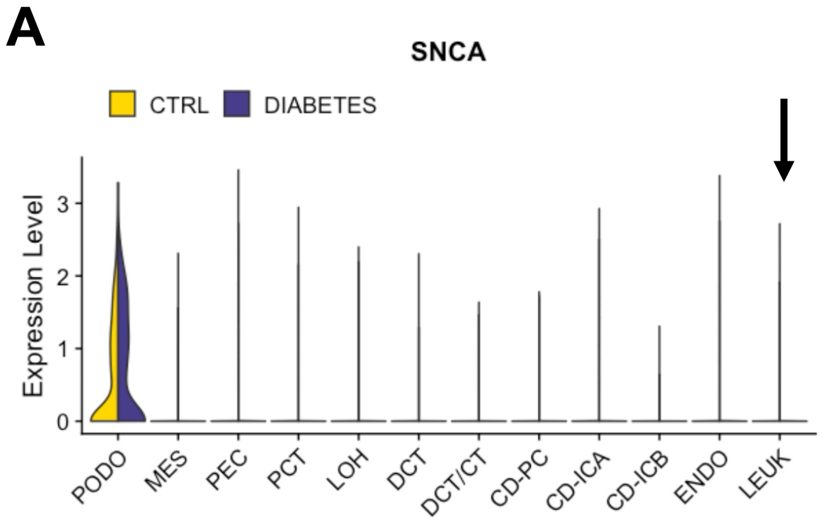


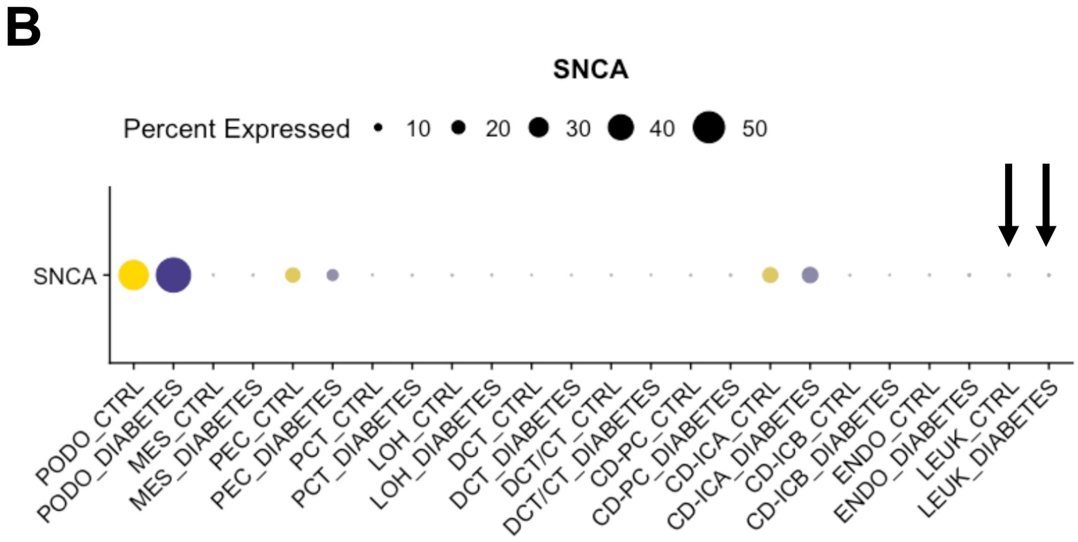

### Supplemental Figure S11

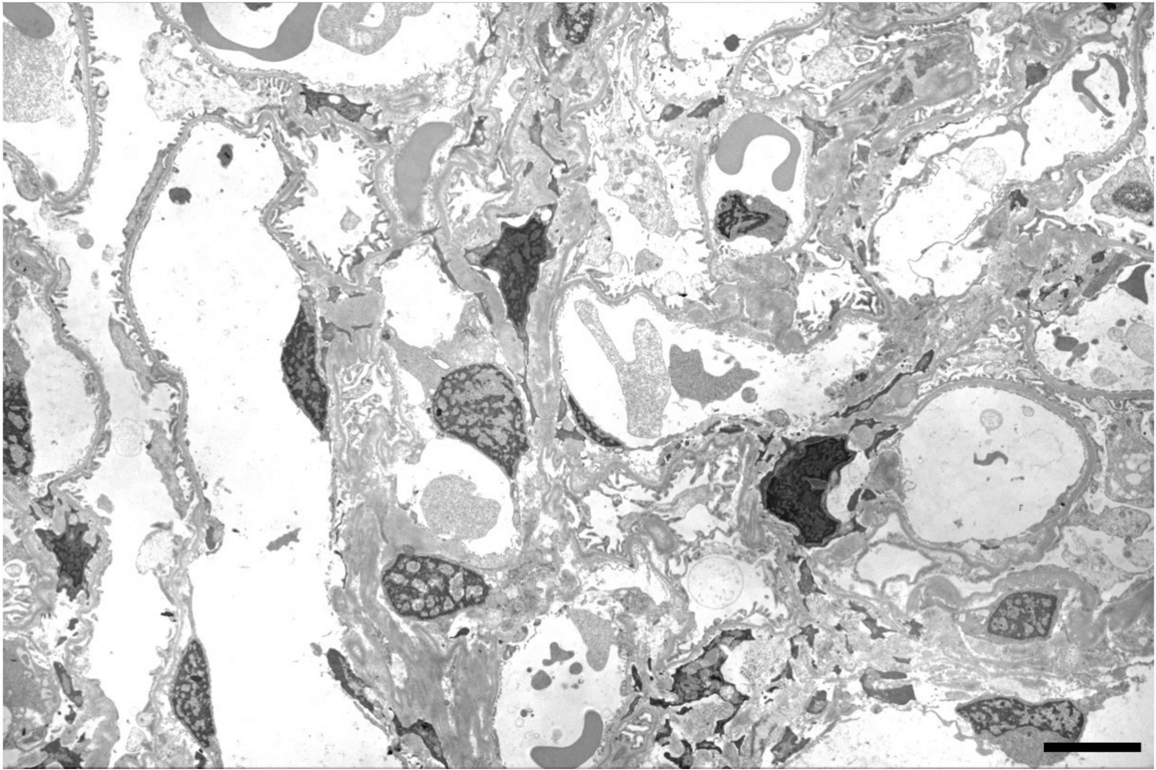
