## Supplemental Figure Legends for "Myeloid FtH Regulates Macrophage Response to Kidney Injury by Modulating Snca and Ferroptosis"

**Figure S1.** (A) Bulk RNA-sequencing data of FtH^fl/fl^ and FtH^Δ/Δ^ kidneys under baseline conditions highlighting genes that encode iron binding proteins (n = 4). Normalized count data of signature genes are plotted. ns = not significant, **P < 0.01. (B) Immunofluorescence staining of Snca in kidneys of FtH^fl/fl^ and FtH^Δ/Δ^ mice under baseline conditions. Scale bar = 50 μm.

**Figure S2.** Immunofluorescence assays were performed using anti-Snca and anti-LY6G antibodies to determine the pattern of expression of these proteins in spleen of FtH^Δ/Δ^ mice. As illustrated neutrophils are not the primary source of Snca expression in FtH^Δ/Δ^ spleens. Scale bar = 100 μm.

**Figure S3.** Expression of Snca in leukocytes as determined by (A) log2 fold change, and (B) transcripts per million reveals monocytes as the major cellular source of expression. Data was generated via accessing publicly available transcriptome analysis platoform. (<https://dice-database.org>)

**Figure S4.** (A) UMAP projection of isolated CD45^+^ (leukocytes) from kidneys of both genotypes, treated with vehicle or AA, passing rigid quality control filtering and after dataset integration, yielding 9 distinct cell clusters (n= 1 per genotype and treatment condition). (B, C) Violin plots showing (B) the number of informative genes per single cell, and (C) the percentage of mitochondrial genes per single cell, all split by batches.

**Figure S5.** (A, B) Heatmap of differentially enriched genes in FtH deficient cells compared to their wildtype counterparts under (A) vehicle, and (B) AA treatment conditions. (C) Dot plots display marker gene expression in isolated leukocytes, comparing FtH-deficient and wildtype monocytes/MΦ treated with vehicle or AA, analyzed six weeks post-final injection to assess ferroptosis-related cell death pathway. Cells are stratified by genotype and treatment condition. Dot size denotes percentage of cells expressing the marker. Color scale represents average gene expression values.

**Figure S6.** (A, B) FtH^fl/fl^ and FtH^Δ/Δ^ kidneys were harvested under baseline conditions and protein expression of (A) HNE, and (B) ACSL4 were examined via Western blotting. (C, D) Densitometric analysis indicates an increasing trend for both (C) HNE, and (D) ACSL4 in baseline FtH^Δ/Δ^ kidneys; however, these results did not achieve statistical significance.

**Figure S7.** (A-C) Three days post-final parenteral iron administration, kidneys, livers, and spleens of FtH^fl/fl^ and FtH^Δ/Δ^ mice were harvested and subjected to a comprehensive metallomics investigation using inductively coupled plasma mass spectrometry (ICP-MS) to measure levels of (A) magnesium, (B) copper, and (C) zinc. ns = not significant, **P < 0.01.

**Figure S8.** Bulk RNA-sequencing data of FtH^fl/fl^ and FtH^Δ/Δ^ kidneys following iron overload highlighting signature genes that encode proteins involved in MΦ iron recycling biological profile (n = 4). Normalized count data of signature genes are plotted. *P < 0.05, **P < 0.01, ***P < 0.001.

**Figure S9.** Reference UMAP, the left panel, shows different annotated cell types. The black dash sequesters cells of monocytic lineage subpopulation of leukocytes and was isolated for better visualization. The middle panel shows expression levels of in healthy kidneys and the right panel demonstrates enrichment of Snca during AKI. Data was generated via accessing kidney precision medicine project website. (<https://www.kpmp.org>)

**Figure S10.** (A) violin plot of Sc-RNAseq demonstrates enrichment of Snca in control and diabetic kidneys that is almost exclusively seen in podocytes. Arrow indicates expression of Snca in isolated leukocytes. (B) Dotplot analysis confirms lack of enrichment of Snca in leukocytes as indicated by arrows. Data was generated via accessing kidney interactive transcriptomics platform developed by Dr. Humphreys. (https://humphreyslab.com/SingleCell/)

**Figure S11.** Representative images of kidney biopsies with an isolated diagnosis of TBM demonstrate thinning of the basement membrane. Scale bar = 5 μm.
