## Supplemental Tables for "Myeloid FtH Regulates Macrophage Response to Kidney Injury by Modulating Snca and Ferroptosis"

| cluster Abbrev | cluster Name | cell Count | Mean Exp | pct Cells Expressing | fold Change | p Val | p Val Adj |
| --- | --- | --- | --- | --- | --- | --- | --- |
| MDC | Monocyte-derived Cell | 1017 | 0.68 | 37.2 | 3.21 | 4.32E-135 | 1.39E-130 |
| POD | Podocyte | 201 | 1.95 | 52.2 | 4.63 | 5.32E-133 | 1.71E-128 |
| IC-A | Intercalated Cell Type A | 1980 | 0.322 | 18.6 | 2.07 | 1.16E-52 | 3.72E-48 |
| MAC-M2 | M2-Macrophage | 465 | 0.852 | 19.6 | 3.43 | 8.51E-42 | 2.74E-37 |
| PEC | Parietal Epithelial Cell | 242 | 0.565 | 21.5 | 2.74 | 2.88E-26 | 9.24E-22 |
| cDC | Classical Dendritic Cell | 298 | 0.647 | 18.1 | 2.95 | 8.44E-23 | 2.71E-18 |
| pDC | Plasmacytoid Dendritic Cell | 36 | 0.666 | 38.9 | 2.99 | 2.39E-22 | 7.69E-18 |
| CNT-IC-A | Connecting Tubule Intercalated Cell Type A | 712 | 0.171 | 14 | 1 | 3.96E-18 | 1.27E-13 |
| cycMNP | Mononuclear Phagocyte (*cycling^2^*) | 14 | 0.59 | 42.9 | 2.92 | 1.39E-13 | 4.46E-09 |
| MON | Monocyte | 459 | 0.223 | 10.5 | 1.39 | 1.06E-07 | 0.00339 |

**Table S1.** Cluster of cells that significantly express Snca mRNA in kidneys of healthy individuals

**Table S2.** Cluster of cells that significantly express Snca mRNA in kidneys of patients with AKI

| cluster Abbrev | cluster Name | cell Count | mean  Exp | pct Cells Expressing | fold Change | p Val | p Val Adj |
| --- | --- | --- | --- | --- | --- | --- | --- |
| PEC | Parietal Epithelial Cell | 178 | 0.858 | 28.7 | 3.88 | 5.69E-73 | 1.83E-68 |
| MAC-M2 | M2-Macrophage | 794 | 0.588 | 21.2 | 3.47 | 5.88E-70 | 1.89E-65 |
| POD | Podocyte | 57 | 1.99 | 47.4 | 5.08 | 7.86E-67 | 2.53E-62 |
| cyc MNP | Mononuclear Phagocyte (*cycling^2^*) | 108 | 0.713 | 25.9 | 3.59 | 5.08E-42 | 1.63E-37 |
| cDC | Classical Dendritic Cell | 1012 | 0.339 | 12 | 2.61 | 8.18E-34 | 2.63E-29 |
| pDC | Plasmacytoid Dendritic Cell | 158 | 0.699 | 20.3 | 3.57 | 8.01E-30 | 2.57E-25 |
| MDC | Monocyte-derived Cell | 582 | 0.449 | 13.4 | 2.99 | 8.58E-29 | 2.76E-24 |

**Table S3.** Characteristics of analyzed kidney tissues

|  | **TBM (n=6)** | **AIN**  **(n=6)** | **ACR (n=6)** |
| --- | --- | --- | --- |
| Age [years] | 46±5.76 | 47±10.79 | 43±6.0 |
| Gender: female [%] | 100% | 50% | 66.6% |
| Max serum creatinine [mg/dL] | 0.9±0.15 | 8.7±4.24 | 4.0±0.76 |

**Table S4.** List of primary antibodies and dilutions used in Immunoblotting, Immunohistochemistry, and Immunofluorescence experiments, related to STAR Methods

| **Primary Antibody** | **Source** | **Catalog Number** | **Dilution for Immunoblotting** |
| --- | --- | --- | --- |
| Purified Mouse Anti-α-Synuclein | BD Biosciences | Cat #610787 | 1:1000 |
| Anti-ACSL4 (FACL4) antibody [EPR8640] | Abcam | Cat #ab155282 | 1:2500 |
| Anti-ferritin heavy chain Antibody (C6) | Santa Cruz Biotechnology | Cat #sc-517438 | 1:10,000 spleens, 1:1000 all other samples |
| Anti-Glyceraldehyde-3-Phosphate Dehydrogenase Antibody, clone 6C5 | Millipore Sigma | Cat #MAB374 | 1:10,000 |
| Anti-4 Hydroxynonenal antibody | Abcam | Cat #ab46545 | 1:2500 |
| Ferritin Light Chain Polyclonal Antibody | Invitrogen | Cat #PA5-19059 | 1:1000 |
| **Primary Antibody** | **Source** | **Catalog Number** | **Dilution for Imunnohistochemistry** |
| α-Synuclein (D37A6) XP® Rabbit mAb | Cell Signaling Technology | Cat #4179S | 1:200 |
| **Primary Antibody** | **Source** | **Catalog Number** | **Dilution for Immunofluorescence** |
| (D37A6) XP® Rabbit mAb | Cell Signaling Technology | Cat #4179S | 1:200 |
| CD11b Monoclonal Antibody (M1/70), Super Bright™ 600, eBioscience™ | ThermoFisher Scientific | Cat #63011282 | 1:200 |
| Alexa Fluor® 700 anti-mouse Ly-6G Antibody | BioLegend | Cat #127622 | 1:200 |

**Table S5.** List of oligonucleotide sequences, related to STAR Methods

| **q-PCR primer sequences** | | | |
| --- | --- | --- | --- |
| **Species** | **Gene Name** | **Primer Name** | **Primer Sequence 5’-> 3’** |
| Mouse | GAPDH | GAPDH FWD | ATC ATC CCT GCA TCC ACT |
| Mouse | GAPDH | GAPDH REV | ATC CAC GAC GGA CAC ATT |
| Mouse | NFE2L2 (NRF2) | NRF2 FWD | CAG CAT AGA GCA GGA CAT GGA G |
| Mouse | NFE2L2 (NRF2) | NRF2 REV | GAA CAG CGG TAG TAT CAG CCA G |
| Mouse | SLC40A1 (ferroportin) | FPN FWD | GTG GAG TAC TTC TTG CTC TGG |
| Mouse | SLC40A1 (ferroportin) | FPN REV | CTG CTT CAG TTC TGA CTC CTC |
| Mouse | SLC7A11 (xCT) | SLC7A11 FWD | GAT TCA TGT CCA CAA GCA CAC |
| Mouse | SLC7A11 (xCT) | SLC7A11 REV | GAG CAT CAC CAT CGT CAG AG |
| Mouse | Spi-C | Spi-C FWD | CCC ACA GAG AAC CCC CTC TA |
| Mouse | Spi-C | Spi-C REV | TGT ACG GAT TGG TGG AAG CC |
| **Genotyping primer sequences** | | | |
| Mouse | Cyclization recombinase (cre) | Cre FWD | GCC AGG CGT TTT CTG AGC ATA C |
| Mouse | Cyclization recombinase (cre) | Cre REV | CAC CAT TGC CCC TGT TTC ACT ATC |
| Mouse | Ferritin heavy chain 1 (FtH1) | FtH1 flox FWD | CCA TCA ACC GCC AGA TCA AC |
| Mouse | Ferritin heavy chain 1 (FtH1) | FtH1 flox REV | CCA TCA ACC GCC AGA TCA AC |
